# SMC motor proteins extrude DNA asymmetrically and contain a direction switch

**DOI:** 10.1101/2023.12.21.572892

**Authors:** Roman Barth, Iain F. Davidson, Jaco van der Torre, Michael Taschner, Stephan Gruber, Jan-Michael Peters, Cees Dekker

## Abstract

Structural Maintenance of Chromosomes (SMC) complexes organize the genome *via* DNA loop extrusion. While some SMCs were reported to do so symmetrically, reeling DNA from both sides into the extruded DNA loop simultaneously, others perform loop extrusion asymmetrically toward one direction only. The mechanism underlying this variability remains unclear. Here, we examine the directionality of DNA loop extrusion by SMCs using *in vitro* single-molecule experiments. We find that cohesin and SMC5/6 do not reel in DNA from both sides, as reported before, but instead extrude DNA asymmetrically, while the direction can switch over time. Asymmetric DNA loop extrusion thus is the shared mechanism across all eukaryotic SMC complexes. For cohesin, direction switches strongly correlate with the turnover of the subunit NIPBL, during which DNA strand switching may occur. STAG1 stabilizes NIPBL on cohesin, preventing NIPBL turnover and direction switches. The findings reveal that SMCs, surprisingly, contain a direction switch subunit.

**Highlights:**

- All eukaryotic SMC complexes extrude DNA asymmetrically.
- Apparent ‘symmetric’ loop extrusion is the result of frequent direction switches.
- n human cohesin, loop-extrusion direction changes require exchange of NIPBL.
- STAG1 stabilizes NIPBL on human cohesin.

## INTRODUCTION

Genomes across the tree of life are organized and constantly reshaped by active processes within the cell. Structural Maintenance of Chromosomes (SMC) protein complexes are important, owing to their major role in chromosome organization^1^. The three major members of the SMC family in eukaryotes are condensin, cohesin, and SMC5/6. All three are capable of manipulating DNA *via* an ATP-dependent process called DNA loop extrusion^2–10^, where loops are created and enlarged stepwise by ATP-binding events to the SMC complex^9,11^. Condensin mainly extrudes DNA loops during mitosis to compact sister chromatids and facilitate chromosome segregation^12,13,13^. Cohesin organizes interphase chromosomes by looping DNA between convergently oriented CCCTC-binding factor (CTCF) proteins^14–17^, giving rise to topologically associated domains in population-average chromosome-conformation-capture^18–22^ as well as imaging experiments^23–25^, which contributes to transcriptional regulation^1,26^ as well as genome integrity^27–29^. SMC5/6 has poorly understood functions in chromosome segregation and genome maintenance^30,31^.

The composition and architecture of the three SMC complexes are highly similar (shown schematically for human cohesin in Figure 1A; reviewed previously^32,33^). Two Smc subunits dimerize at their hinge domain and are connected via coiled-coil arms to their heads. The heads harbour ATP-binding domains related to those of ATP-binding cassette (ABC) transporters, which dimerize upon ATP binding to catalyse its hydrolysis. A flexible kleisin subunit bridges the Smc ATPase heads and provides binding sites for two HAWK (HEAT protein associated with a kleisin) proteins in cohesin and condensin, or KITE (kleisin interacting tandem winged-helix element) proteins in SMC5/6.

**Figure 1:**
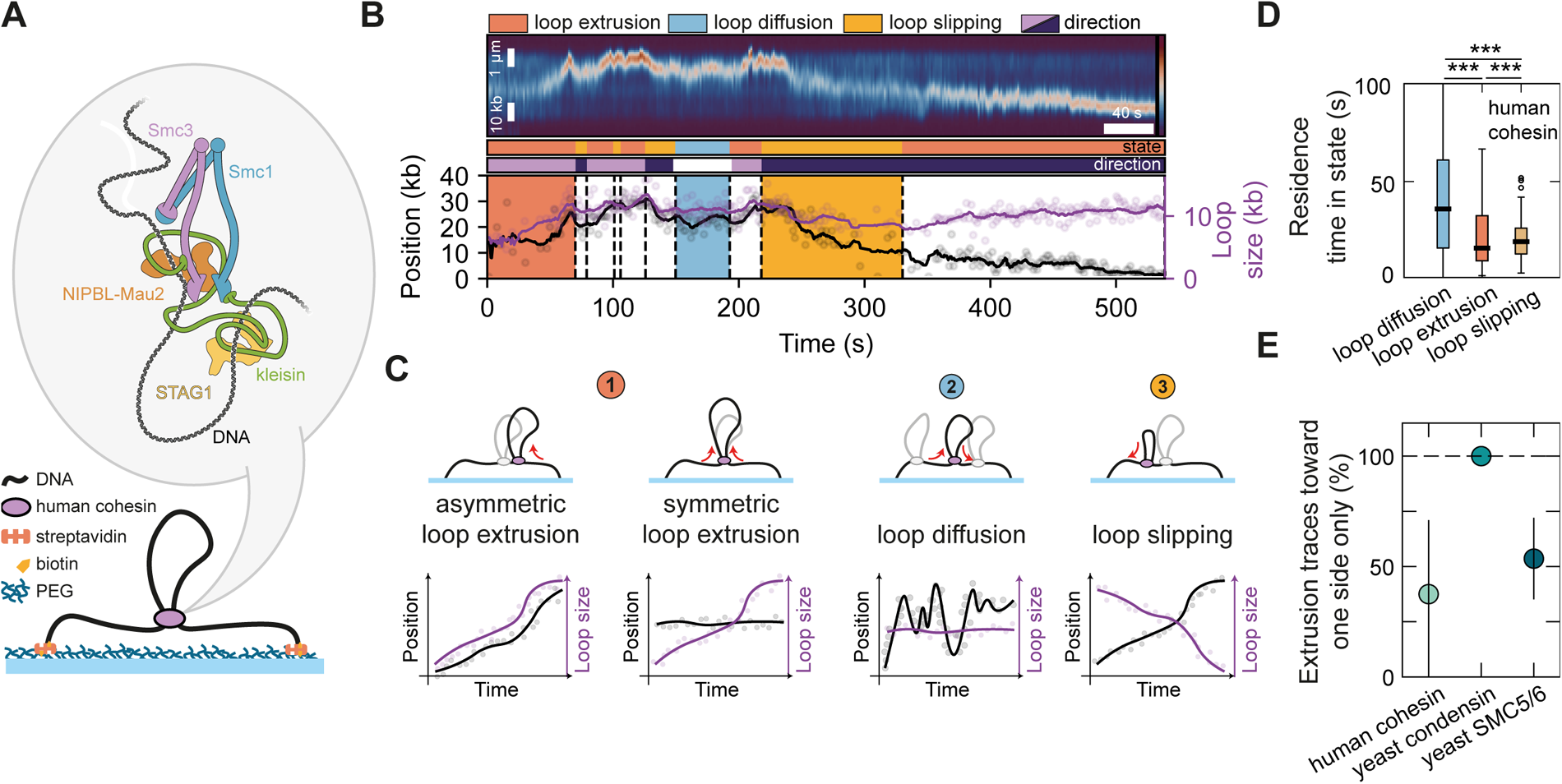
SMC-mediated DNA loop extrusion is interspersed with loop-diffusion and loop-slipping events. **(A)** Illustration of the DNA loop extrusion assay and the architecture of human cohesin containing Smc3, Smc1, Scc1 (kleisin), NIPBL-Mau2, and STAG1/2 (here only STAG1 is used). The corresponding subunit names for yeast condensin are Smc4, Smc2, Brn1, Ycs4, and Ycg1. The corresponding subunit names for yeast SMC5/6 are Smc5, Smc6, and Nse4 with additional Nse subunits Nse1, Nse2, and Nse3. **(B)** A typical kymograph of cohesin-mediated DNA loop extrusion. Loop position and size are quantified. Their time propagation allows to discern phases of loop extrusion, diffusion and slipping. Extrusion and slipping have a direction given by the local sign of the derivative of the loop position with respect to time. **(C)** Illustration of loop position and size propagation in time for asymmetric and symmetric extrusion, loop diffusion, and slipping. **(D)** Duration of diffusion, extrusion, and slipping states (N = 93, 344, 257, respectively) for human cohesin. Black horizontal lines are median values, the box extends between the first and third quartile, and the whiskers extend to 1.5*IQR. Statistical significance was assessed by a Mann-Whitney test with Bonferroni correction. (***: p<0.001). **(E)** Fraction of SMC complexes with at least two extrusion events that displayed only unidirectional extrusion (N = 13, 49, 33 for cohesin, condensin, and SMC5/6, respectively). Error bars denote the binomial 95% confidence interval. See also Figures S1 and S2.

Despite the overall highly conserved architecture of eukaryotic SMCs and their common ability to extrude DNA loops^32,34^, they appear to significantly differ in their loop extrusion directionality. Here, we use the following terminology to describe the directionality: asymmetric loop extruders incorporate DNA only from one side into the loop, while symmetric extruders reel in DNA from both sides simultaneously into the loop. Unidirectional extruders undergo sequential phases of asymmetric extrusion (possibly interrupted by pauses) that always occur in the same direction, whereas bidirectional extruders exhibit phases of asymmetric extrusion but here the side from which DNA is incorporated into the loop switches direction over time. Yeast condensin is an asymmetric and unidirectional extruder, that is, it reels DNA into the loop strictly from one side^2,35^, while human cohesin was reported to reel DNA from both sides into the loop and was thus considered to be a symmetric DNA loop extruder^3,4^. Dimeric yeast SMC5/6 similarly was reported to be a symmetric extruder^7^. Given the structural similarity of these complexes^8,32,33^, the reason for these differences is unclear. This variability undermines whether the loop extrusion mechanism^36^ is shared among all SMC complexes.

Here, we experimentally measured the directionality of DNA loop extrusion by human cohesin, yeast condensin, and yeast SMC5/6 at a single-molecule level. Close inspection of their loop extrusion dynamics revealed that cohesin and SMC5/6 undergo short phases of active extrusion, characterized by loop growth, which are interspersed by diffusion^3^ of the loop and SMC across the DNA, and loop slipping, the gradual loss of the previously extruded loop. Focusing on the extrusion phases, we find that all three monomeric SMC complexes extrude DNA strictly asymmetrically within these phases. While the condensin holocomplex is a strictly unidirectional extruder, cohesin and monomeric SMC5/6 undergo frequent direction switches, making them bidirectional loop extruders. Furthermore, we found that human cohesin undergoes direction switches upon exchange of its HEAT subunit NIPBL. STAG1, in contrast, stabilizes NIPBL on cohesin, impeding NIPBL turnover, and thereby prevents extrusion direction switches. These findings indicate that the DNA loop extrusion mechanism is inherently asymmetric and is likely common to all SMC complexes.

## RESULTS

### Phases of active DNA loop extrusion are interspersed by loop diffusion and loop slipping

To characterize the dynamics of DNA loop extrusion by human cohesin in detail, we reconstituted DNA loop extrusion with purified proteins at the single-molecule level *in vitro.* To this end, λ-DNA, which served as a substrate for loop extrusion, was tethered at both ends onto a polyethylene glycol-passivated glass surface at an extension of ∼30% of its contour length. DNA was visualized using SYTOX Orange intercalating dye, and imaged using highly inclined and laminated optical sheet (HILO) microscopy (Methods). Human cohesin and ATP were subsequently introduced into the flow channel and the flow was stopped to record cohesin-mediated loop extrusion in the absence of buffer flow (Figure 1A). Kymographs were constructed that served as the basis for quantifying the size of the loop and its position (Figure 1B, Methods).

The time propagation of these two independent quantities, loop size and position, defines distinct phases into which loop extrusion traces were segmented (Figure 1C): (i) Phases of active DNA loop extrusion exhibited an increase in loop size (Figures 1C and red shading in Figure 1B), (ii) extended ‘loop diffusion’ periods in which the loop size remained approximately constant, while the loop position changed in a diffusive manner (Figures 1C, blue shading in Figure 1B, and Figure S3A), and (iii) gradual shrinking of loops (Figures 1B, yellow shading in Figure 1C, and Figure S3B), as noted previously as a possible outcome of cohesin-CTCF encounters^37^. Phases of loop stalling, e.g. by reaching the end of surface-tethered DNA molecules, were excluded (e.g. Figure 2). This gradual shrinkage is distinct from a sudden step-wise loop rupture, which resolves the loop within one or few frames (∼100 ms) ^2,35^. Loop shrinkage typically led to only a partial (i.e. incomplete) loss (Figures S1A, S1B, S1G, and S3B-E). Quantitatively, cohesin remained longest in the diffusive state (41 ± 44 s, median ± SD), while spending 16 ± 17 s (median ± SD) in active extrusion, which was comparable to the time spent in the loop slipping phases (20 ± 15 s, median ± SD; Figure 1D). We tested whether even shorter phases existed, which could have been missed by acquiring images at a frame rate of 5-10 frames per second (fps). Acquired images at 50 fps yielded no significantly shorter phases however (Methods, Figures S1C-S1F).

**Figure 2:**
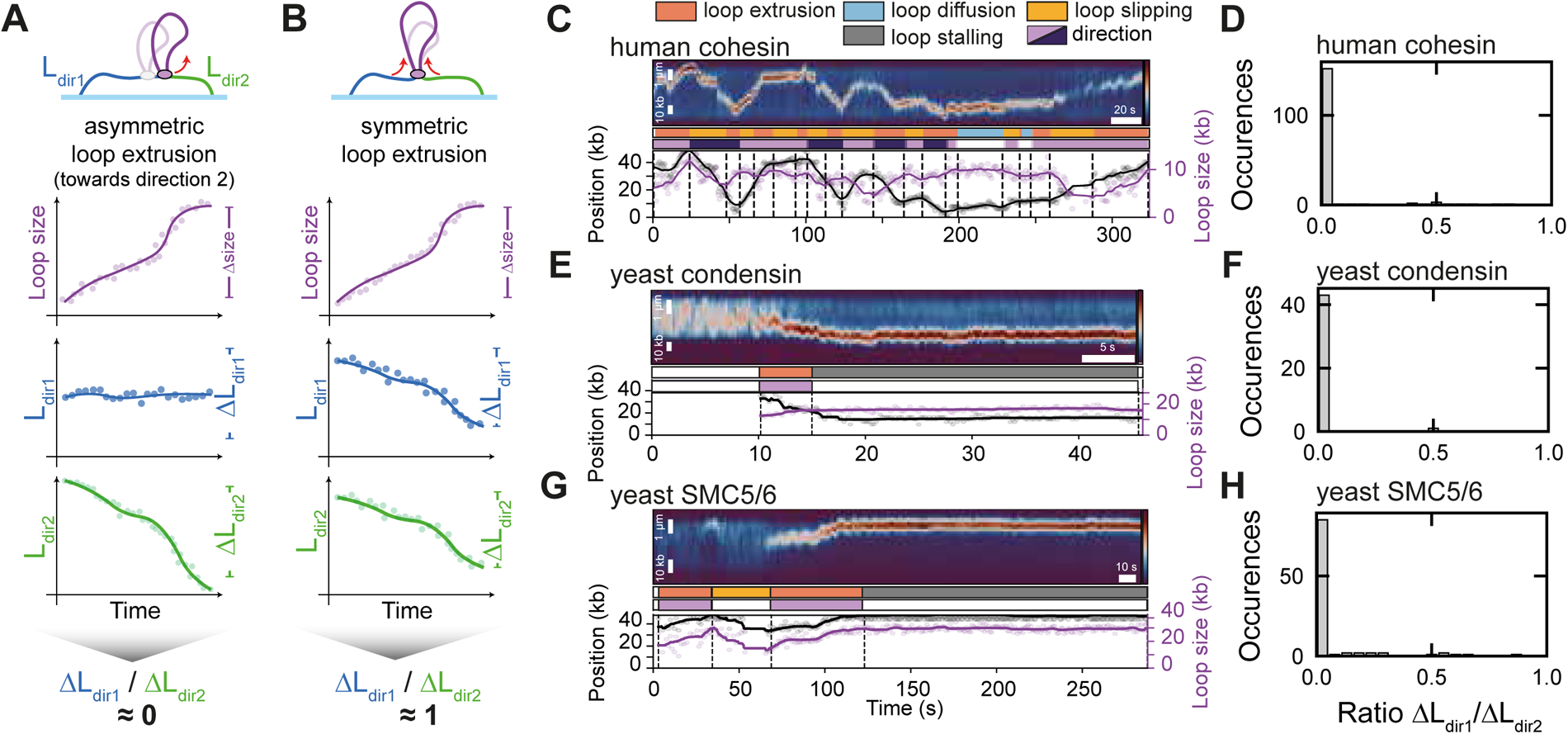
All eukaryotic SMC complexes extrude DNA in an asymmetric manner. **(A)** Illustration of loop size and amount of DNA left and right of the loop in time for asymmetric loop extrusion. **(B)** Illustration of loop size and amount of DNA left and right of the loop in time for symmetric loop extrusion. The ratio of changes in amount of DNA left and right of the loop is discriminative between asymmetric and symmetric extrusion. **(C)** Exemplary loop extrusion kymograph of cohesin with loop position and size quantification and segmentation into phases of extrusion, diffusion, and slipping. **(D)** Distribution of the ratio ΔL_dir1_/ΔL_dir2_ for extrusion events of human cohesin (N = 166 events). **(E)** Exemplary kymograph of unidirectional DNA loop extrusion for yeast condensin. **(F)** Distribution of the ratio ΔL_dir1_/ΔL_dir2_ distribution for yeast condensin (N = 44 events). **(G)** Exemplary kymograph of unidirectional DNA loop extrusion for yeast SMC5/6. **(H)** Distribution of the ratio ΔL_dir1_/ΔL_dir2_ distribution for yeast SMC5/6 (N = 100 events). See also Figures S1 and S2.

To examine how the loop extrusion dynamics among the three eukaryotic SMC complexes differed, we repeated the experiments and analyses with budding yeast condensin and budding yeast SMC5/6 (Methods, Figure S2). First, we tested whether the purified SMC5/6 used here is able to perform DNA loop extrusion, as reported previously^7^ (Figure S2). Indeed, we confirmed that SMC5/6 exhibits loop extrusion of DNA with similar characteristics as reported (Figures S2B-S2E), although we found that yeast SMC5/6 – like human cohesin and yeast condensin – extrudes already as a monomer (Figures S2F-S2I), while Pradhan et al.^7^ reported loop extrusion only for SMC5/6 dimers. This held true both for SMC5/6 expressed and purified from *E.coli* (ref. 38) as well as from *S. cerevisiae* (ref. 7; compare Figures S2G and S2I). While SMC5/6 exhibited dynamics comparable to cohesin, condensin exhibited only brief phases of diffusion (7 ± 4 s, median ± SD) while extrusion phases lasted 13 ± 9 s (median ± SD), and loop slipping was hardly ever observed (1/65 phases; Figure S1J).

### All eukaryotic SMC complexes extrude DNA asymmetrically

The growth and concomitant change in the loop position is a proxy for the direction of DNA loop extrusion, as illustrated in Figure 1C. By monitoring the direction of travel of the loop extrusion phases during the experiment (from the start of the extrusion until loop dissolution or the end of acquisition), we measured the fraction of SMC complexes that showed multiple extrusion phases in the same or opposite directions. While condensin is a strictly unidirectional extruder (all extrusion phases moved toward one side of the DNA), 40-60% of cohesin and SMC5/6 complexes that show at least two extrusion phases extruded toward both sides during the course of the experiment (Figure 1E).

Cohesin-mediated loop extrusion was mostly interspersed with either a diffusion or a slipping phase (or both) between subsequent extrusion phases (Figure S1I); switches between extrusion and slipping phases were most common (Figure S1J). Direction changes without intervening diffusion or slipping phases were less common, as only approximately 25% of direction changes occur during loop extrusion, i.e. without observable intermittent diffusion or slipping phase (Figure S1I). The sequence of extrusion, diffusion, and slipping phases can be described as a first-order Markov chain, that is, the transition probabilities between the states solely depend on the current state, indicating that the system is memory-less (Figure S3A and S3B).

Asymmetric and symmetric DNA loop extruders are characterized by their ability to reel DNA from one or both sides into the loop, respectively^2,7^. For an asymmetric extruder, the stretch of DNA on one side of the SMC becomes progressively shorter, while the length of the other side remains constant^2^ (Figure 2A). In contrast, both arms become simultaneously shorter as the loop is being enlarged by a symmetric extruder (Figure 2B). Thus, the ratio of the length changes on both sides of the loop during loop extrusion constitutes a quantity that we name an ‘symmetry indicator’ which is close to 0 for an asymmetric extruder (since ΔL_dir1_ ≈ 0 in Figure 2A) and ≍1 for a symmetric extruder. To determine whether cohesin extrudes DNA asymmetrically or symmetrically, we computed the symmetry indicator Δ*L_dir1_*/Δ*_dir2_* within the segmented loop extrusion phases. We found that 92 ± 4 % (mean ± 95% binomial confidence interval, N = 166) of all extrusion phases exhibited a value of Δ*L_dir1_*/Δ*L_dir2_* below 0.1 (Figure 2D). Extrusion phases of yeast condensin, a known asymmetric extruder^2^ (Figure 2E), used here as a positive control, exhibited a similar distribution with 98 ± 4 % (mean ± 95% binomial confidence interval, N = 44, Figure 2F) of all values of Δ*L_dir1_*/Δ*L_dir2_* below 0.1. Monomeric yeast SMC5/6 (Figure 2G) also exhibited asymmetric extrusion phases (86 ± 7 % [mean ± 95% binomial confidence interval] of Δ*L*_*dir1*_/Δ*L_dir2_* values below 0.1, N = 100, Figures 2H and S1L). Notably, the slipping phases of cohesin and SMC5/6 were also asymmetric (Figure S1M and S1N), suggesting that loop slipping is the result of weak DNA binding at either the motor or anchor side of the SMC complexes, but not both.

We conclude that all eukaryotic SMC complexes extrude DNA asymmetrically, strictly toward one direction at a time; however, cohesin and SMC5/6 may switch the direction of extrusion between two successive extrusion phases.

### NIPBL excess increases the frequency of direction switches in cohesin-mediated loop extrusion

Even though yeast condensin is a strictly unidirectional DNA loop extruder, the complex extrudes towards both directions alternatingly when Ycg1, condensin’s ‘anchor’ (ref. ^35^), is deleted, or when the strong DNA binding site within the safety belt^39^, is mutated. These results suggest that weakening or dissolution of at least one DNA-binding site is required to permit direction switches during DNA loop extrusion. Human cohesin requires a reservoir of unbound NIPBL-Mau2 for ongoing DNA loop extrusion and loop maintenance^3^ and *in vivo* FRAP experiments indicated that NIPBL-Mau2 hops from cohesin to cohesin in cells^40^, suggesting that NIPBL-Mau2 is only transiently bound to cohesin as NIPBL-Mau2 heterodimers continuously exchange on cohesin. As NIPBL-Mau2 contains a strong DNA-binding site^41^, we hypothesized that dissociation of NIPBL-Mau2 from the complex weakens the DNA-binding strength close to the Smc3 subunit, which could subsequently lead to extrusion direction switches.

To test this, we performed cohesin-mediated DNA loop extrusion experiments with varying concentrations of NIPBL-Mau2 to modulate the average occupancy of NIPBL-Mau2 on cohesin. In loop extrusion experiments where NIPBL-Mau2 was added at a sub-stoichiometric ratio to cohesin, mostly unidirectional extrusion was observed (Figure 3A, ratio of NIPBL-Mau2 to cohesin = 0.1, and Figure 3D). Strikingly, upon increasing the NIPBL-Mau2:cohesin ratio, more traces with extrusion phases toward both directions were observed (Figures 3A and 3E-3G). In addition, the fraction of cohesin complexes that underwent diffusion and loop slipping increased with increasing NIPBL-Mau2:cohesin concentration ratios (Figure S3C and S3D). Bidirectional extrusion cannot be attributed to the dimerization of cohesin, as suggested previously^4^, since in both uni- and bidirectional extrusions, mostly a single cohesin complex colocalized with the loop^3^ (Figure S3E-S3G). The increase in bi-versus unidirectionally extruding cohesin complexes with increasing NIPBL-Mau2:cohesin ratio did not coincide with an enhanced direction exchange frequency (Figure 3B). Instead, the loop lifetime, the time between the beginning of the first extrusion phase until loop dissolution, increased as a larger excess of NIPBL-Mau2 over cohesin was supplied (Figure 3C), in line with previous loop maintenance assays^3^. Subsequent extrusion phases were indistinguishable in terms of loop extrusion rate (Figure S3H). These results suggest that rebinding of NIPBL to cohesin stabilizes loops and that cohesin can undergo additional loop extrusion cycles in any direction upon rebinding of a NIPBL molecule.

**Figure 3:**
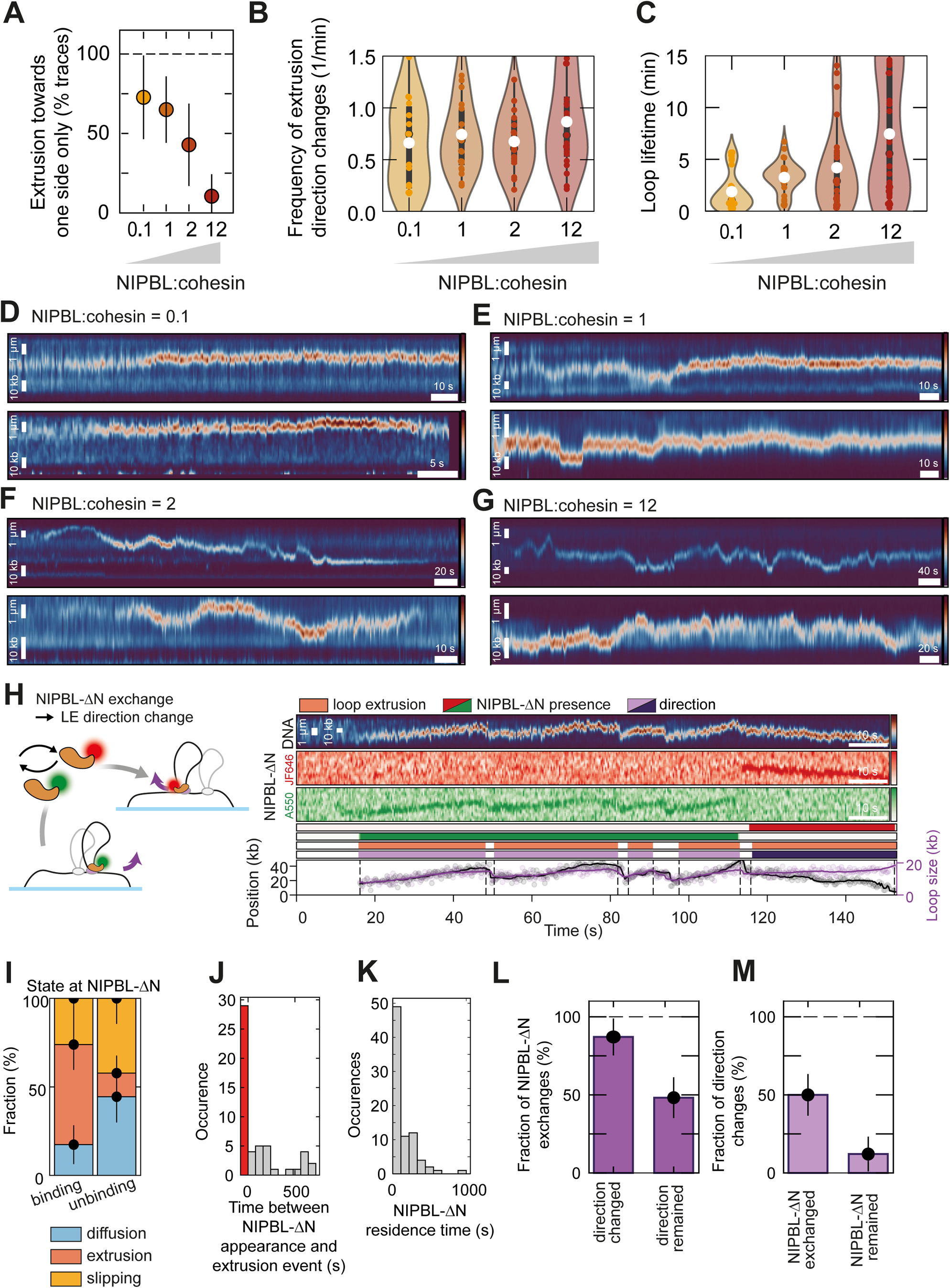
DNA loop extrusion direction changes coincide with an exchange of NIPBL-ΔN. **(A)** Fraction of cohesin complexes undergoing at least two extrusion events that displayed only unidirectional extrusion for varying ratios of NIPBL-Mau2:cohesin at 50 mM NaCl without additional STAG1 (N = 41, 30, 17, 26 from left to right). Error bars denote the binomial 95% confidence interval. **(B)** Frequency of extrusion direction changes for varying ratios of NIPBL-Mau2:cohesin at 50 mM NaCl (N = 41, 30, 17, 26). The white dot denotes the mean. The box shows the quartiles of the dataset and the whiskers extend to 1.5*IQR. **(C)** Loop lifetime between the first extrusion event and loop disruption for varying ratios of NIPBL-Mau2:cohesin at 50 mM NaCl (N = 41, 31, 25, 37) without additional STAG1. The white dot denotes the mean. The box shows the quartiles of the dataset and the whiskers extend to 1.5*IQR. **(D)** Exemplary kymographs of cohesin-mediated loop extrusion at NIPBL-Mau2:cohesin = 0.1 at 50 mM NaCl. **(E)** Exemplary kymographs of cohesin-mediated loop extrusion at NIPBL-Mau2:cohesin = 1 at 50 mM NaCl. **(F)** Exemplary kymographs of cohesin-mediated loop extrusion at NIPBL-Mau2:cohesin = 2 at 50 mM NaCl. **(G)** Exemplary kymographs of cohesin-mediated loop extrusion at NIPBL-Mau2:cohesin = 12 at 50 mM NaCl. **(H)** Exemplary trace where the exchange of NIPBL-ΔN coincides with a change in the loop extrusion direction. Cohesin extrudes toward the lower end as long as the ATTO 550-labeled NIPBL-ΔN is bound to cohesin. After exchange to a JF646-labeled NIPBL-ΔN, cohesin extrudes toward the upper end. HeLa cohesin was used with NIPBL-ΔN-ATTO550 and NIPBL-ΔN-JF646. **(I)** Probability of states at which NIPBL-ΔN binds to cohesin (N = 46) and at which NIPBL-ΔN dissociates (N = 45). Binding and unbinding at extrusion phases was only observed concomitant with the start/end of extrusion phases, respectively. The bar height indicates the probability of a given state and the error bar denotes the binomial 95% confidence interval. **(J)** Distribution of time between NIPBL-ΔN appearance and the next extrusion event (N = 56). The red bar represents the occurrences when NIPBL-ΔN appearance coincided with the start of an extrusion event. **(K)** Distribution of NIPBL-ΔN residence times (N = 47). **(L)** Fraction of cohesin molecules for which NIPBL-ΔN exchanged in between direction changes (87 ± 12 %, N = 31) and the fraction of cohesin molecules on which NIPBL-ΔN exchanged without direction change between successive extrusion phases (48 ± 13 %, N = 56) at 50 mM NaCl and in the absence of additional STAG1. Error bars denote the binomial 95% confidence interval. **(M)** Fraction of direction changes between successive extrusion phases during which NIPBL-ΔN exchanged (50 ± 13 %, N = 54) and the fraction of direction changes between successive extrusion phases during which NIPBL-ΔN remained (12 ± 11 %, N = 33) at 50 mM NaCl and in the absence of additional STAG1. Error bars denote the binomial 95% confidence interval. See also Figures S3 and S4.

### In the absence of NIPBL, loops can diffuse and slip but cannot be extruded

Next, we directly visualized whether an exchange of NIPBL-Mau2 on cohesin occurs. For this, we expressed and purified an N-terminal truncation of NIPBL (NIPBL-ΔN, ref. ^3,41^) which was split into two batches that were labelled differentially with ATTO 550 (NIPBL-ΔN-A550) and Janelia Fluor 646 (NIPBL-ΔN-JF646) fluorophores (Methods, Figure S4A). We then performed cohesin-mediated DNA loop extrusion experiments with an equimolar ratio of NIPBL-ΔN-A550 and NIPBL-ΔN-JF646 and monitored the presence and absence of NIPBL-ΔN in the two colors colocalizing with the loop (Figures S4B-E and Figure 3H). Cohesin did not show any extrusion phases in the absence of NIPBL-ΔN. NIPBL-ΔN binding to cohesin typically coincided with the start of a loop-extrusion phase, but NIPBL also bound to cohesin during diffusion and slipping phases (Figure 3I, left bar). In the latter case, the next extrusion phase followed within a few minutes (142 ± 222 s [mean ± SD], Figure 3J). Dissociation of NIPBL-ΔN occurred most often during diffusion or slipping phases, on average 31 ± 53 s (mean ± SD) after the last extrusion phase (Figure S4F), and only rarely at the end of extrusion phases (Figure 3I, right bar) after NIPBL-ΔN spent on average 161 ± 186 s (mean ± SD) bound to cohesin (Figure 3K). Cohesin without bound NIPBL-ΔN was typically rebound by NIPBL-ΔN after ∼1-2 min (Figure S4G) though cohesin without bound NIPBL-ΔN was observed for up to 5 min (at NIPBL-ΔN:cohesin ratio of 0.1-1 in these experiments), which is in agreement with the report that in the absence of NIPBL-Mau2 most loops dissociated within 8 min (ref. 3). The diffusion constant of diffusing loops as well as the loop slipping rate were not significantly different in the presence or absence of NIPBL-ΔN (Figures S4H and S4I), suggesting that NIPBL does not contribute to DNA binding during the diffusion and slipping phases. Because NIPBL is known to contribute to DNA clamping onto the ATPase heads upon ATP binding^42,43^, this suggests that diffusion and slipping phases represent an ATP-unbound states of cohesin.

### Extrusion direction switches coincide with an exchange of NIPBL on cohesin

Next, we analysed how an exchange of NIPBL-ΔN relates to the extrusion direction switching of cohesin. We considered an exchange of NIPBL-ΔN if the fluorescence intensity of one label was replaced with an intensity of the other label, but also when the fluorescence intensity of one label disappeared and re-appeared at least 2 frames later. Note that imposing this restriction allowed us to monitor exchanges between NIPBL-ΔN carrying the same label only as long as cohesin remained unbound by NIPBL-ΔN for at least 600 ms. Figure 3H shows an exemplary kymograph of cohesin-mediated loop extrusion, in which first (15 – 112 s) four successive extrusion phases move toward the upper end of the DNA molecule with intermittent loop slipping phases. During all these unidirectional phases, a NIPBL-ΔN-A550 molecule colocalized with the DNA loop (presumably bound to cohesin^3^), which however dissociated from cohesin at t = 113 s. At t = 116 s, a NIPBL-ΔN-JF646 bound to the cohesin, which coincided with a direction switch as extrusion of the loop now proceeded toward the lower end of the DNA molecule. We monitored such an appearance and disappearance of NIPBL-ΔN in relation to direction switches for multiple molecules (additional examples in Figure S4B-S4E). In 86 ± 12 % of the cases in which a direction change was observed, there was also an exchange of NIPBL-ΔN between extrusion phases in opposite directions (e.g. Figure S4B). When subsequent extrusion phases occurred in the same direction, an exchange of NIPBL-ΔN was observed in 52 ± 12 % of the cases (Figure 3L, example in Figure S4D). Conversely, an exchange of NIPBL-ΔN coincided in 47 ± 12 % of the cases with a direction switch, while only 14 ± 11 % of the cases in which a direction switch was observed did not correlate with an exchange of NIPBL-ΔN (Figure 3M).

These data show that extrusion direction switches coincide an exchange of NIPBL, strongly suggesting that NIPBL exchange is a necessary step to permit extrusion direction switches. The opposite is however not true: an exchange of NIPBL does not necessarily yield a direction switch. It is possible that NIPBL molecules carrying the same fluorophore did exchange within a time period that could not be resolved (using the time resolution of 300 ms used in these experiments). Events in which direction changes were observed without concomitant NIPBL exchange might have been caused by such events (Figures 3L, first bar and 3M, second bar).

### STAG1 is dispensable for cohesin-mediated DNA loop extrusion at low ionic strength

Cohesin is an asymmetric extruder (Figure 2) that can switch extrusion directions (Figure 3). A working model that allows asymmetric extruders to switch directions is the ‘DNA strand exchange model’ in which DNA temporarily unbinds from the extruder’s motor and anchor side and switches between them^6,35^. In line with such a DNA strand exchange model, yeast condensin is unable to perform bidirectional extrusion as long as DNA is tightly bound at Ycg1, condensin’s anchor side^35^. By contrast, human cohesin is able to extrude in a bidirectional manner, despite the fact that the cohesin purification contains STAG1, cohesin’s Ycg1 analogue. This can be explained by three possibilities that are not mutually exclusive. First, DNA-binding at the STAG1-kleisin DNA binding site might be weaker than that Ycg1-kleisin (‘safety belt’)^39,41^ because no safety belt has yet been identified in cohesin^3,32,33^. This could result in temporary DNA unbinding from STAG1-kleisin. Second, STAG1, like NIPBL-Mau2, could dissociate from cohesin and exchange with a STAG1 molecule from solution. However, since loops can be maintained in the absence of STAG1 during buffer flow, this exchange would have to occur either at slow rates or STAG1 is not essential to maintain DNA loops, in line with the observations that condensin-ΔYcg1 (ref. ^35^) can extrude loops. And third, STAG1 could be, like Ycg1 (ref. ^35^), facultative for DNA loop extrusion by cohesin or only needed for loop initiation in order to overcome the bending energy associated with bending DNA into a small loop^44^.

To test the latter, we first performed cohesin-mediated DNA loop extrusion experiments using single-chain cohesin, which is purified in the absence of STAG1 from *Sf9* insect cells (Figure S3I, ref. 3) using a constant 2.5-fold excess of NIPPBL-Mau2 over cohesin. The single-chain cohesin trimer with NIPBL-Mau2 was largely unable to initiate DNA loop extrusion in the absence of STAG1 at 50 mM NaCl (Figure S4J, ref. 3). However, lowering the ionic strength to 25 mM NaCl permitted DNA loop extrusion without STAG1 (Figure S4J), in line with a ΔYcg1 version of *Chaetomium thermophilum* condensin which was also able to extrude DNA (bidirectionally)^35^. Thus, DNA binding at STAG1 helps to initiate DNA loops, but STAG1 is not strictly necessary at low ionic strength. In these conditions, the single-chain cohesin trimer with NIPBL-Mau2 also switched extrusion directions (Figures 4A and 4B) and was able to maintain loops similar to recombinant cohesin (purified with STAG1) at 50 mM NaCl (Figure 4C).

**Figure 4:**
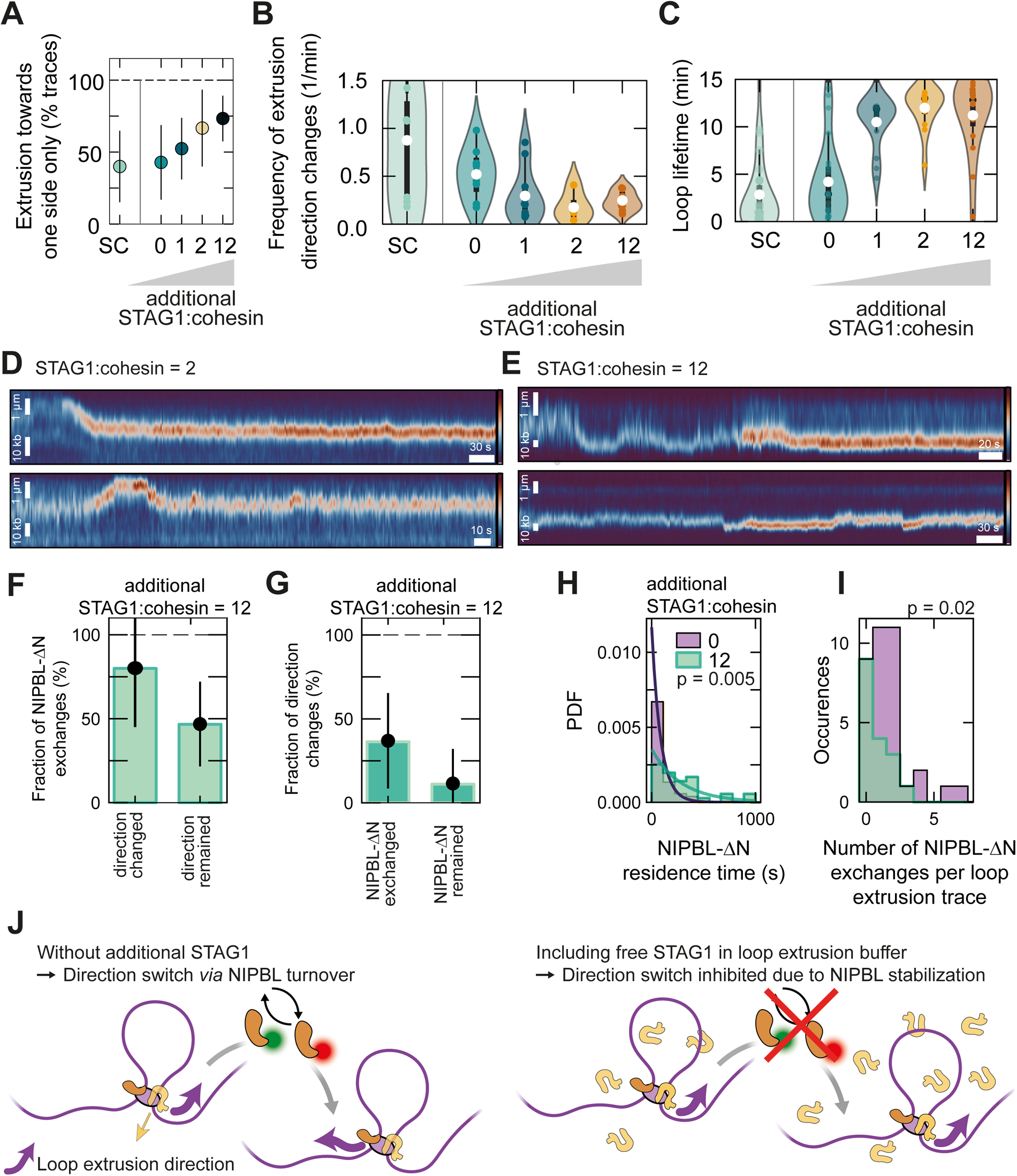
STAG1 stabilizes NIPBL on cohesin and prevents direction changes. **(A)** Fraction of cohesin complexes undergoing at least two extrusion events that displayed only unidirectional extrusion for single-chain cohesin in the absence of STAG1 (SC; loop extrusion at 25 mM NaCl) and increasing ratio of STAG1:cohesin (recombinant cohesin) at a constant ratio of NIPBL-Mau2:cohesin = 2.5 at 50 mM NaCl (N = 15, 14, 21, 12, 30 from left to right). The white dot denotes the median. The box shows the quartiles of the dataset and the whiskers extend to 1.5*IQR. **(B)** Frequency of extrusion direction changes for single-chain cohesin in the absence of STAG1 (SC; loop extrusion at 25 mM NaCl) and increasing ratio of STAG1:cohesin at NIPBL-Mau2:cohesin = 2.5 at 50 mM NaCl (N = 7, 8, 10, 4, 8). The white dot denotes the mean. The box shows the quartiles of the dataset and the whiskers extend to 1.5*IQR. **(C)** Loop lifetime between the first extrusion event and loop disruption for single-chain cohesin (SC) in the absence of STAG1 (loop extrusion at 25 mM NaCl) and increasing ratio of STAG1:cohesin at NIPBL-Mau2:cohesin=2.5 at 50 mM NaCl (N = 25, 25, 16, 13, 16). The white dot denotes the mean. The box shows the quartiles of the dataset and the whiskers extend to 1.5*IQR. **(D)** Exemplary kymographs of cohesin-mediated loop extrusion at NIPBL-Mau2:cohesin = 2.5 at 50 mM NaCl and a 2-fold excess of free STAG1 added to the loop extrusion buffer. **(E)** Exemplary kymographs of cohesin-mediated loop extrusion at NIPBL-Mau2:cohesin = 2.5 at 50 mM NaCl and a 12-fold excess of free STAG1 added to the loop extrusion buffer. **(F-G)** As for Figures 3L and 3M but in the presence of 12-fold molar excess of STAG1 over cohesin (NIPBL-ΔN:cohesin = 2.5). The distribution does not change with respect to the condition without additional STAG1, suggesting that exchanges of NIPBL-ΔN determine extrusion direction switches also in the presence of additional STAG1 (N = 5, 15 in panel F; N = 11, 9 in panel G from left to right. **(H)** as for Figure 3K comparing the NIPBL-ΔN residence time in the absence (purple) and presence (green; N = 32) of a 12-fold molar excess of STAG1 over cohesin. Data in the absence of additional STAG1 is the same as shown in (I). Statistical significance was assessed by a Mann-Whitney U rank test on two independent samples. **(I)** Comparison of the number of NIPBL-ΔN exchanges in the absence (purple; N = 36) and presence (green; N = 17) of a 12-fold molar excess of STAG1 over cohesin. Statistical significance was assessed by a Mann-Whitney U rank test on two independent samples. **(J)** Illustration of the relationship between direction changes, NIPBL turnover, and NIPBL stabilization of STAG1 by NIPBL. In the absence of additional STAG1 in the DNA loop extrusion buffer, NIPBL turnover is required to exchange the loop extrusion direction. Likely, STAG1 dissociates from cohesin after loop initiation. If free STAG1 is included in the loop extrusion buffer, direction switches are inhibited by stabilization of NIBPL on cohesin, presumably due to an interaction between NIPBL and STAG1. See also Figure S4.

### STAG1 stabilizes NIPBL on cohesin

To test whether STAG1 may exchange on cohesin analogously to NIPBL, we performed loop extrusion experiments with recombinant cohesin (containing STAG1) and additionally added an increasing amount of purified STAG1 to cohesin at 50 mM NaCl. An excess of STAG1 in loop extrusion reactions yielded an increasing amount of unidirectional cohesin complexes (Figures 4A and compare examples in Figures 3D-3G and Figures 4D-4E)., i.e., an excess of STAG1 has the opposite effect of NIPBL on the loop extrusion directionality (cf. Figure 3A). In contrast to NIPBL, an excess of STAG1 lowered the frequency of extrusion direction changes about 2-fold (Figure 4B) but it also increased the loop lifetime (Figure 4C). As cohesin complexes in the absence of STAG1 are unable to mediate DNA loop extrusion at 50 mM NaCl (Figure S4J and ref. 3), the observed loops must have been initiated from cohesin-NIPBL-STAG1 complexes, however it is unclear whether STAG1 remained stably bound to cohesin after the start of loop extrusion or whether STAG1 may exchange on cohesin.

An exchange of NIPBL permits extrusion direction switches (Figure 3L). Given that an excess of STAG1 promotes unidirectional extrusion, it is unclear which one of the HEAT subunits, NIPBL or STAG1, controls direction switches. To test whether NIPBL turnover alone is predictive for extrusion direction switches, or whether STAG1 directly contributes to direction switches, we monitored extrusion direction switches and the exchange of NIPBL-ΔN-A550/JF646 in DNA loop extrusion experiments with a 12-fold excess of STAG1 over cohesin. We observed that the fraction of NIPBL-ΔN exchanges during which the extrusion direction was maintained in the presence of additional STAG1 was indistinguishable from experiments without additional STAG1 (compare the right bars in Figures 3L and 4F [p = 0.78, two-tailed Fisher’s exact test], as well as the left bars in Figures 3M and 4G [p = 0.75, two-tailed Fisher’s exact test]).

We thus conclude that direction switches solely coincide with the exchange of the NIPBL subunit. However, how does STAG1 impact cohesin’s loop extrusion directionality? Since the frequency of extrusion direction changes is independent of the NIPBL concentration (Figure 3B) and direction changes require NIPBL dissociation (Figure 3L), the dissociation rate of NIPBL from cohesin is independent of the NIPBL concentration. However, the frequency of extrusion direction changes depended on the excess of STAG1 over cohesin. This suggests that the dissociation rate of NIPBL from cohesin is dependent on STAG1. To test this prediction, we measured the residence time of NIPBL-ΔN-A550/JF646 on cohesin during DNA loop extrusion. Indeed, we found that NIPBL-ΔN resides on average 1.7-times longer on cohesin in the presence of a 12-fold excess of STAG1 over cohesin compared to experiments without additional STAG1 (278 ± 239 s at STAG1:cohesin=12 [mean ± SD], p = 0.005 from a Mann-Whitney U rank test on two independent samples, Figure 4H). We also observed a 2.1-fold lower number of NIPBL-ΔN exchanges on cohesin per loop extrusion trace (0.8 ± 0.9 at 12-fold excess of STAG1 compared to 1.7 ± 1.6 without additional STAG1 [mean ± SD], p = 0.02 from a Mann-Whitney U rank test on two independent samples, Figure 4I). We conclude that cohesin can change its extrusion direction *via* the turnover of NIPBL while that binding of STAG1 to the cohesin complex stabilizes NIPBL, thus preventing direction switches (Figure 4J).

## DISCUSSION

### DNA loop extrusion is conserved, asymmetric, and can switch direction

DNA loop extrusion is an important molecular mechanism across the tree of life. While yeast condensin has been reported to extrude DNA from one side only^2^, human cohesin^3,4^ and yeast SMC5/6 (ref. 7) were found to reel in DNA from both sides, suggesting that cohesin and SMC5/6 are symmetric extruders. These differential loop extrusion dynamics questioned whether the DNA loop extrusion mechanism is conserved among Eukaryotic SMC complexes. Using single molecule *in vitro* reconstitution of DNA loop extrusion by all three Eukaryotic SMCs and quantification of their loop extrusion dynamics, we set out to clarify whether an asymmetric DNA loop extrusion mechanism may constitute the common *modus operandi* of the Eukaryotic SMCs. Indeed, we found that all monomeric Eukaryotic SMCs extrude DNA strictly asymmetrically (Figure 2). However, cohesin and SMC5/6 are able to switch the direction of extrusion (Figure 1E). These results strongly suggest that the DNA loop extrusion mechanism is shared among the eukaryotic SMCs (potentially including the prokaryotic SMCs as well), is inherently asymmetric, and that the DNA loop extrusion cycle permits direction switches.

Our results suggest that direction switches require an exchange of NIPBL on human cohesin (Figure 3). Strikingly, the propensity of cohesin and condensin to switch extrusion directions can be tuned by modulating the ability to bind DNA at either the HEAT-A subunit (NIPBL/Ycs4, respectively), HEAT-B subunit (STAG1/Ycg1), or both. For cohesin, dissociation of NIPBL likely weakens a DNA binding site close to Smc3 (the motor side) because NIPBL forms a strong DNA-binding site on the cohesin holocomplex^41^. Not only NIPBL but also STAG1 may be able to dissociate from cohesin during loop extrusion which would weaken DNA binding at cohesin’s anchor side (Figure 4). This is in line with previous data showing that condensin is able to extrude DNA loops bidirectionally in the absence of its analogous HEAT-B subunit Ycg1 (ref. ^35^). Note that the exchange of DNA-binding subunits is an extreme measure to temporarily weaken DNA binding at these subunits. In order to allow direction switches *via* DNA strand exchange (see below), DNA would have to temporarily dissociate from Ycs4. The fact that direction switches are observed for ΔYcg1 condensin, even though Ycs4 was intact ^35^, suggests that DNA can indeed temporarily dissociate from Ycs4 (and potentially other HEAT and KITE subunits) without complete subunit dissociation from the complex. This might also explain why both uni-as well as bidirectional extrusion has been observed concomitantly for human condensin I and II (ref. 10) and for *X. laevis* condensin and cohesin from egg extracts^5^, where no subunit turnover is known. The activity of human cohesin, but not yeast condensin, is regulated *in vivo* by the replacement of NIPBL by Pds5/WAPL and by the balance of STAG1- and STAG2-bound cohesin^45^. This might explain why subunits (at least NIPBL) are dynamically exchanged on cohesin, while they are not for other SMC proteins.

### Mechanistic models that enable DNA strand exchange

How can a DNA-loop-extrusion model account for direction switches? We envision two possible scenarios that extend the recently proposed reel-and-seal model^36^, based on previous considerations in which DNA strands between the motor and anchor sides are exchanged (‘DNA strand exchange’) to enable extrusion direction switches^6,35^, see Figures 5 and S5. The stem of the DNA loop is bound to least two DNA binding sides: the motor subunit (NIPBL) and the anchor subunit (STAG1/2). Weak DNA binding at the STAG1-kleisin interface or STAG1 dissociation could hand over the previously bound DNA strand to a binding site on the Smc1 ATPase head^41^ since STAG1 and Smc1 can be positioned in close spatial proximity. Indeed, Ycg1 was found in close proximity to both ATPase heads in an ATP-bound state (in the absence of DNA)^46^ and a structure prediction by AlphaFold2 (ref. ^47^) suggests that STAG1 may be bound to the Smc1 but not to the Smc3 ATPase head (Figures S5C and S5D). In the first variant, dissociation of NIPBL could similarly transfer DNA to the DNA-binding site on the Smc3 ATPase head^41^ in the ATP-unbound state (Figure 5 upper arc and Figure S5G). The DNA strands can then exchange between the Smc subunits, potentially exploiting the spatial proximity induced by ATPase head engagement upon ATP binding. Rebinding of NIPBL to cohesin selects the Smc3-bound DNA strand as the strand which is subsequently extruded into the loop.

**Figure 5:**
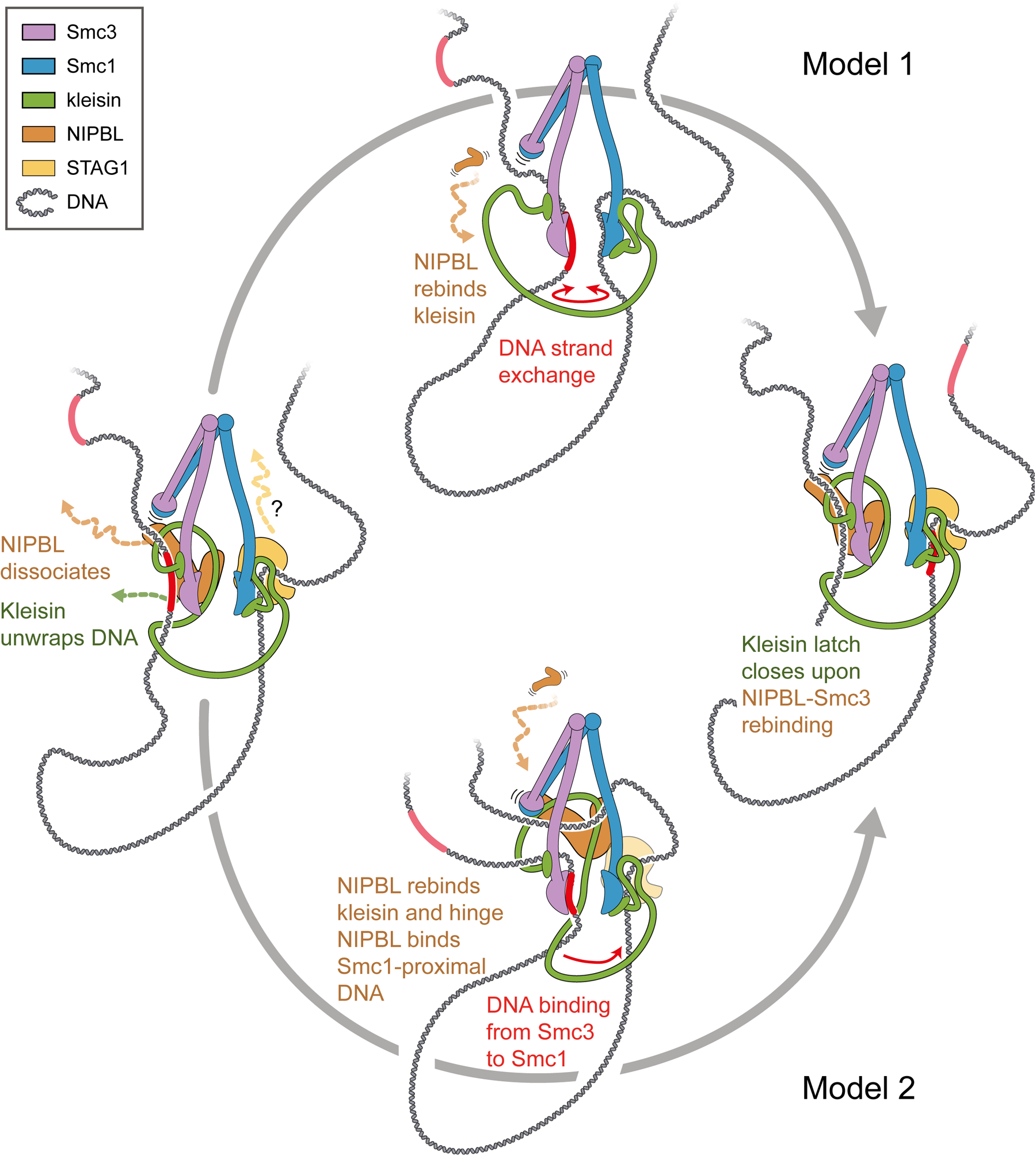
Potential pathways of loop extrusion direction change via DNA strand exchange mediated by NIPBL exchange. Upon dissociation of NIPBL, the kleisin unwraps DNA (left panel). To prevent loss of the Smc3-proximal DNA (red), the Smc3 ATPase head binds DNA. STAG1 may dissociate from cohesin concomitantly, while the Smc1 ATPase head can bind the Smc1-proximal strand (faint DNA strand). DNA strand exchange may occur spontaneously between the ATPase heads (upper panel). Upon rebinding of NIPBL to kleisin (upper panel) and closing of the kleisin latch, the former Smc1-proximal strand is poised for the next loop extrusion cycle (right panel). Alternatively, NIPBL may re-bind the cohesin complex by binding to kleisin and the Smc hinge (lower panel). The NIPBL-kleisin-hinge complex may bind either DNA strand which is the one on which loop extrusion subsequently proceeds. If the former Smc1-proximal DNA strand is chosen, the Smc3-proximal DNA strand exchanges to the DNA binding site at the Smc1 ATPase head or at STAG1. The binding of NIPBL-kleisin to the Smc3 ATPase head prepares the complex for the next loop extrusion cycle. See also Figure S5.

Instead of DNA strand exchange based on the DNA binding sites on both ATPase heads, a second scenario is that NIPBL could associate with the hinge upon rebinding^41^. In this stage, the NIPBL-hinge complex could bind either the Smc3- or Smc1-proximal DNA strand, poising each one of them to be extruded into the loop in the subsequent extrusion step (Figure S5H). Both possibilities, DNA strand exchange *via* the DNA binding sites on the ATPase heads or *via* NIPBL-hinge association may or may not lead to DNA strand exchange and require DNA to transiently dissociate from NIPBL and STAG1. This is in line with the observation that NIPBL turnover can, but does not necessarily lead to direction switching. Strong DNA binding at either HEAT-A or HEAT-B would inhibit these DNA-strand-exchange pathways, explaining why yeast condensin does not undergo direction switches in the presence of the strong safety belt^2,2,39^. Note that both pathways do not necessarily require subunit exchange, as DNA unbinding from both subunits (and for model 2 unbinding of NIPBL from Smc3 and binding to the hinge) is sufficient. The models are also sufficiently general to be applicable for direction changes observed for human condensin I and II (ref. 10) as well as for *X. laevis* condensin and cohesin from egg extracts^5^. It is unclear whether ATP binding or hydrolysis contribute to the conformational changes required to enable DNA strand exchange. Since direction changes occur much less frequently than loop extrusion steps, the potential transient states involved in DNA strand exchange will be difficult to resolve using structural methods like cryo-EM.

An excess of STAG1 during DNA loop extrusion stabilizes NIPBL on cohesin (Figure 4H and 4I). An experimentally solved structure of cohesin suggests that NIPBL is sandwiched between the engaged ATPase heads and STAG1 (ref. ^42^) which could prevent NIPBL dissociation in the ATP-bound state (Figure S5A), a conformation which was also observed in yeast condensin^8^. Fission yeast Psc3^Scc3^ and Ssl3^Scc2^ were also found to interact in isolation^48^ and protein structure prediction by AlphaFold2 furthermore suggest a direct interaction between NIPBL and STAG1 (Figure S5B and ref. ^49^). No experimentally solved structure including the Smc ATPase heads, NIPBL, and STAG1 in the apo state exists to date. We thus turned to AlphaFold2 to predict the position of NIPBL and STAG1 on the respective Smc subunits. The predictions suggest that STAG1 is bound to Smc1 but not to Smc3 in the absence of NIPBL (Figures S5C and S5D). When NIPBL is included, NIPBL is placed in between Smc3 and STAG1 (Figure S5E) and on Smc1 (Figure S5F). This and the finding that Ssl3^Scc2^ and Psc3^Scc3^ interact in the absence of Smc subunits and DNA (Figure S5B, ref. ^48,49^) opens up the possibility that the direct association between NIPBL and STAG1 prevents NIPBL turnover as long as STAG1 is part of the cohesin complex, even in the absence of ATP.

### The physiological relevance of extrusion direction switching

In human and yeast cells, NIPBL is present at a slightly sub-stoichiometric ratio to Scc1 (ref. ^50–53)^. However, Pds5 is more abundant than NIPBL which could potentially result in an excess of NIPBL over Pds5-unbound cohesin *in vivo.* Based on this stoichiometry, we thus expect that NIPBL turnover and direction switching occur *in vivo,* in line with studies which found that Scc2 is not stably associated with chromosomal cohesin^54^ and hops between cohesin complexes in HeLa cells^40^. We anticipate that the direction changes described here are essential for proper genome organization by SMC complexes^55^. Simulations have shown that unidirectional extruders are unable to recapitulate the experimentally observed interphase chromosome organization, in particular the formation of TADs, even if the anchor site is only diffusively bound to DNA^56^, while asymmetric extrusion with Z-loop formation and direction switching could recapitulate mitotic chromosome formation^56,57^. In contrast, asymmetric extrusion with switching adequately reproduces the interphase chromosome organization. These simulations suggested that cohesin has to undergo direction switches approximately once per minute, yielding in the order of 10 switches before cohesin dissociation from DNA. This estimate is in fair agreement with our *in vitro* results (Figures 3B and 4B). Further support that direction switching by cohesin is likely to occur *in vivo* comes from the observation of extrusion “fountains” (in zebrafish^58^, in *C. elegans*^59^, “plumes” in mouse cells^60^, and “jets” in mouse thymocytes^61^), regions of enhanced contact frequency in Hi-C maps, that emanate from a narrow region at which cohesin is preferentially loaded and broadening as extrusion proceeds away. Polymer simulations suggest that extrusion direction switching can explain the formation of these fountains^58^. In the simulations, the authors supposed that direction switching resulted from collisions between SMC proteins, but the fact that SMCs can traverse each other^62^ makes this unlikely. Instead, direction switching is likely inherent to cohesin-mediated DNA loop extrusion.

### Limitations of the study

In experiments which probe the turnover of NIPBL-ΔN, exchanges between NIPBL molecules carrying the same fluorophore faster than 600 ms (two time points) may appear like stably bound complexes. This might potentially account for that fact that not all direction switches were accompanied by a NIPBL exchange in these experiments (Figure 3L) and that direction switches were observed also within periods in which no NIPBL exchange was observed (Figure 3M). Early experiments demonstrating DNA loop extrusion by human cohesin were performed under sideflow^3^. In these experiments, both arms of the non-extruded DNA shrank simultaneously. The results obtained in the present manuscript, in the absence of continuous buffer flow, suggest that this symmetric extrusion is the result of rapid direction switches with concomitant NIPBL turnover. To check whether buffer flow might accelerate direction switching, we performed sideflow experiments as in Davidson *et al.* (ref. ^3^) to probe whether short extrusion phases could be observed that alternatingly proceed along one arm or the other. We failed, within the imaging noise, to observe such periods at a time resolution of 100 ms. Furthermore, we attempted to visualize a potential (fast) NIPBL turnover on cohesin during sideflow experiments. However, due to a propensity of NIPBL to accumulate (which is to some extent alleviated by using NIPBL-ΔN), we observed only an increase of fluorescence owing to NIPBL at the loop stem and on DNA in general, preventing us from observing an exchange of single molecules.

Furthermore, the present study provides evidence that STAG1 stabilizes NIPBL on cohesin. However, whether this entails an exchange of STAG1 on cohesin during cohesin-mediated DNA loop extrusion remains unclear. Unequivocal proof of STAG1 turnover would require purified and differentially labelled STAG1 to perform exchange experiments as for NIPBL (Figure 4).

Finally, we showed that monomeric SMC5/6 can extrude DNA loops asymmetrically. The fact that monomeric yeast SMC5/6 purified from *E.coli* as well as from yeast cells is able to extrude DNA loops is in contrast to a previous study which showed that only SMC5/6 dimers are able to extrude DNA^7^. We also observed that SMC5/6 is able to switch directions. However, our study focused on human cohesin, leaving the molecular mechanism underlying direction changes of SMC5/6 unresolved.

## EXPERIMENTAL MODEL AND SUBJECT DETAILS

### Protein expression and purification

Human cohesin, as well as NIPBL-Mau2, were expressed in and purified from *Sf9* insect cells and HeLa cells as described previously^3^. Recombinant cohesin from *Sf9* insect cells was fluorescently labelled with JF646 as described previously^3^. *S. cerevisiae* condensin was expressed in and purified from *S. cerevisiae* as described previously^2^. *S. cerevisiae* SMC5/6 was expressed, purified, and fluorescently labelled from *E.coli* as described previously^9^. *S. cerevisiae* SMC5/6 was expressed, purified, and fluorescently labelled from *S. cerevisiae* as described in Pradhan *et al.* (ref. ^7^) with small modifications. The cells were lysed and bound to IgG Sepharose as described in Pradhan *et al.* (ref. ^7^). Then the beads were washed in STO500 buffer (50 mM Tris-HCl pH 8.0, 500 mM NaCl, 2 mM MgCl_2_, 0.5 mM tris(2-carboxyethyl)phosphine (TCEP), 10% glycerol, 0.1% IGEPAL CA-630) and eluted in 3 ml of STO500 containing TEV protease.

NIPBL-ΔN was expressed and purified as described previously^3,41^ for NIPBL-MAU2 except that before elution from FLAG-M2 agarose, FLAG-M2 agarose beads were resuspended to one bead volume in 25 mM NaH_2_PO_4_/Na_2_HPO_4_ pH 7.5, 150 mM NaCl, 5 % glycerol, 50 mM imidazole pH 7.5 and split in half for fluorescent labelling with Atto550 and JF646 (see below).

## METHOD DETAILS

### Fluorescent labelling of SMC5/6 and estimation of the labelling efficiency

SMC5/6 hexamer purified from *E.coli* with a C-terminal HALO tag on Nse2 was incubated with a 2-fold molar excess of Janelia-Fluor646 HaloTag Ligand (Promega; 12 µM Smc5/6 hexamer + 24 µM label). After incubation for 1 h at 25°C, the reaction was stopped and excess label was removed using a Zeba Spin desalting column (Thermo Fisher) equilibrated in SEC buffer (20 mM Tris pH 7.5, 250 mM NaCl, 1 mM TCEP).

SMC5/6 purified from *E.coli* and JF646 concentrations were measured independently using absorption measurements at 280 nm for SMC5/6 and at 646 nm for JF646, yielding a fraction of 0.58 ± 0.11 (N = 8 independent measurements) JF646 molecules per SMC5/6 hexamer. Fluorescence Correlation Spectroscopy (FCS) was performed on samples of 10 nM SMC5/6 hexamers in a buffer containing 40 mM Tris-HCl pH 7.5, 50 mM NaCl, 2.5 mM MgCl_2_, 1 mM TCEP at room temperature. The resulting autocorrelation functions was well described by a fit with a single component, indicating that no free fluorophores were present.

SMC5/6 purified from yeast (roughly 1 µM concentration) was incubated with 20 µM of Alexa647-SNAP label overnight at 4°C in the presence of 1 mM DTT. The complex was then purified using a Zeba Spin column equilibrated in STO500 buffer. The labelling efficiency of SMC5/6 purified from yeast was estimated similarly and resulted in a labelling efficiency of 0.22 ± 0.02 (N = 3 independent measurements).

### Fluorescent labelling of NIPBL-ΔN

Janelia Fluor 646 SE (Tocris; 6148) was dissolved in dimethylformamide (DMF) at 20 mM. HaloTag Amine (O2) Ligand (Promega; P6711) was dissolved in dimethylformamide (DMF) at 50 mM. HaloTag Amine (O2) Ligand was added to JF646 SE in a 248.7 µl reaction in DMF while stirring and was then supplemented with 1.3 µl diisopropylethylamine (Sigma) (final concentrations: 7 mM JF646 SE, 2.3 mM HaloTag ligand, 31 mM diisopropylethylamine). Labelling reactions were incubated at room temperature for ∼ 16 hours protected from light. The reaction mixture was diluted 10-fold in solvent A (5 % acetonitrile, 0.1 % formic acid) and purified in five successive reverse phase HPLC runs using an Ultimate 3000 (ThermoFisher Scientific) equipped with a Kinetex 5μ XB-C18 100A, 250 x 4.6mm column using the following gradient of solvents: A (5 % acetonitrile, 0.1 % formic acid), B (acetonitrile, 0.1 % formic acid); 0 – 100 % B in 30 min at a flow rate of 0.8 ml / min. Peak fractions were pooled, lyophilized, resuspended in DMSO, frozen in liquid nitrogen and stored at -80°C. Fractions were analysed by incubating with purified HaloTag protein for 15 min at room temperature followed by SDS-PAGE and excitation using a Bio-Rad ChemiDoc MP.

Atto550 NHS-ester (ATTO-TEC; AD 550-35) was dissolved in dimethylformamide (DMF) at 20 mM. HaloTag Amine (O2) Ligand was added to Atto550 NHS-ester in a 245 µl reaction in DMF while stirring and was then supplemented with 5 µl diisopropylethylamine (Sigma) (final concentrations: 25 mM Atto550 NHS-ester, 8.3 mM HaloTag ligand, 115 mM diisopropylethylamine). Labelling reactions were performed as for Janelia Fluor 646 SE, except that reaction mixture was diluted 10-fold in solvent A (40% acetonitrile, 0.1 % Trifluoroacetic acid) and reverse phase HPLC was performed using the following gradient of solvents: A (40% acetonitrile, 0.1 % Trifluoroacetic acid), B (acetonitrile, 0.1 % Trifluoroacetic acid); 0 – 100 % B in 30 min at a flow rate of 0.8 ml / min.

To label recombinant NIPBL-ΔN with Atto550-HaloTag Ligand and JF646-HaloTag Ligand, the purification was split in half. Each aliquot was supplemented with excess Atto550-HaloTag Ligand or JF646-HaloTag Ligand and incubated for 15 min at room temperature protected from light. Beads were washed extensively with recombinant cohesin purification buffer 3 and bound protein was eluted and concentrated as described for NIPBL-MAU2 in ^3^.

### Double-tethered DNA assay for single-molecule imaging of DNA loop extrusion

DNA loop extrusion experiments and imaging were performed essentially as described previously^2,15,62–64^ with one round of pegylation with 5 mg/ml methoxy-PEG-N-hydroxysuccinimide (MW 3500, LaysanBio) and 0.05 mg/ml biotin-PEG-N-hydroxysuccinimide (MW 3400, LaysanBio) in 0.1 M NaHCO_3_, 0.55 M K_2_SO_4_ buffer^65^. Subsequently, three more rounds of pegylation with MS(PEG)_4_ (MW 333.33, Thermo Fisher) in the same buffer were applied. Slides and coverslips were incubated overnight at 4°C in the dark and then washed with ultrapure water and dried by a N_2_ stream.

After assembly of flow chambers by sandwiching coverslip and drilled glass slide using double-sided tape and sealing the edges using epoxy glue, 0.5 mg/ml streptavidin was incubated for 1 min in loop extrusion buffer (see below) omitting glucose oxidase, intercalating dyes, ATP, and protein. The flow cell was then washed with 100 μl of the same buffer. End-biotinylated λ-DNA (48.5 kbp, ref. ^2^) was then introduced at a flow speed to reach an end-to-end extension of about 6-7 μm (30% of the contour length). Unbound DNA was washed out with up to 200 μl buffer. The DNA was found to be nicked, preventing a buildup of plectonemic supercoils during DNA loop extrusion^63,66–68^.

DNA Loop extrusion reactions by human cohesin were carried out in a buffer containing 40 mM Tris-HCl pH 7.5, 50 mM NaCl (unless otherwise denoted), 2.5 mM MgCl_2_, 2.5% D-glucose, 2 mM Trolox, 10 nM catalase, 18.75 nM glucose oxidase, 100 nM Sytox Orange, 0.5 mg/ml BSA, 1 mM DTT, 1 mM ATP with 10-100 pM cohesin (from Sf9 insect cells) and the indicated concentration of NIPBL-Mau2 at 37°C. For experiments in which cohesin-JF646 was photobleached, glucose oxidase was omitted from the loop extrusion buffer. For experiments with differentially labelled NIPBL-ΔN, the buffer was adjusted to 25 mM NaCl and 25 nM Sytox Green. Part of the experiments were performed with cohesin purified from HeLa cells in which case 1 nM HeLa cohesin was used and part was performed with cohesin purified from *Sf9* cells. The two cohesin purifications behaved similarly and the results of these experiments were pooled.

DNA loop extrusion experiments with yeast condensin were carried out in a buffer containing 40 mM Tris-HCl pH 7.5, 50 mM NaCl, 2.5 mM MgCl_2_, 2.5% D-glucose, 2 mM Trolox, 10 nM catalase, 18.75 nM glucose oxidase, 100 nM Sytox Orange, 0.5 mg/ml BSA, 1 mM DTT, 1 mM ATP with 1 nM cohesin at room temperature (22 °C).

DNA loop extrusion experiments with yeast SMC5/6 from *E.coli* were carried out in a buffer containing 40 mM Tris-HCl pH 7.5, 50 mM NaCl, 2.5 mM MgCl_2_, 2.5% D-glucose, 2 mM Trolox, 10 nM catalase, 18.75 nM glucose oxidase, 100 nM Sytox Orange, 0.5 mg/ml BSA, 1 mM TCEP, 1 mM ATP with 3 nM SMC5/6 hexamer at 30°C or in the same buffer but with 100 mM NaCl, 7.5 mM MgCl_2_ as before^9^. DNA loop extrusion experiments with yeast SMC5/6 from yeast were performed in the same buffer with 100 mM NaCl, 7.5 mM MgCl_2_ as in Pradhan *et al.* (ref. ^7^).

All data were acquired using an exposure time of 100 ms per channel unless otherwise stated. For DNA loop extrusion without monitoring differentially labelled NIPBL-ΔN, the 561 nm-channel was used to image Sytox Orange-stained DNA and the 640 nm-channel was used to alternatingly monitor the SMC complex. The effective time resolution in these experiments was thus 200 ms. For experiments with differentially labelled NIPBL-ΔN, the 480 nm channel was used to image Sytox Green-stained DNA, the 561 nm channel was used to image NIPBL-ΔN-A550, and the 640 nm channel was used to image NIPBL-ΔN-JF646. The time resolution in these experiments was thus 300 ms. Experiments with the aim to visualize exchanges of NIPBL-Mau2 were performed with the N-terminal truncation of NIPBL, NIPBL-ΔN-A550 and NIPBL-ΔN-JF646, because NIPBL-ΔN has a smaller propensity to accumulate/aggregate during DNA loop extrusion experiments and was thus more suitable to image single NIPBL molecules.

To evaluate whether imaging at 200 ms time resolution potentially omits short phases of loop extrusion, diffusion, or slipping, experiments with human cohesin were also performed while imaging Sytox Orange-stained DNA at 20 ms exposure time, yielding thus a 10-fold higher sampling rate compared to the remainder of the experiments.

Image series were acquired for 15 min after flush-in of SMC proteins.

### Fluorescence Correlation Spectroscopy (FCS)

Fluorophore diffusion measurements were conducted using version 1 coverslips on a Picoquant Microtime 200 microscope, which was operated using Symphotime software at room temperature. To focus a 640 nm laser, a 60x Olympus UPSLAPO 60XW water immersion objective with a working distance of 280 µm and a numerical aperture of 1.2 was used. Prior to the experiment, the molecular brightness of a Alexa647 fluorophore solution was optimized by adjusting the correction collar of the objective. The emission light was subsequently directed through a 50 µm pinhole, split by a dichroic mirror, and filtered through a 600/75 optical band pass filter (Chroma, Bellow Falls). Fluorescence emission was collected by single-photon avalanche-diode detectors (PD5CTC and PD1CTC, Micro Photon Devices, Bolzano).

For fitting of the FCS curves, the size of the confocal volume was determined from measurements of the free Alexa647 by fitting a single-component diffusion model with triplet state (a diffusion constant of 297 μm^2^/s was used^69^). The axial and lateral sizes of the confocal volume were fixed for further analysis. FCS amplitudes and diffusion coefficients were subsequently fitted for curves recorded on SMC5/6-JF646 containing samples.

### Mass photometry experiments

Coverslips (Menzel, 24x60mm, #1.5) were washed three times alternatingly with isopropanol and milli-Q water, dried in a nitrogen stream and mounted onto a Refeyn One^MP^ device (Refeyn). Data was acquired using the AcquireMP software (Refeyn Ltd.) and analysed using DiscoveryMP (Refeyn Ltd.). Before the measurement, a mass calibration was recorded to allow conversion between contrast to mass using a protein standard (Invitrogen NativeMark™ Unstained Protein Standard) with masses of 1236, 1048, 720, 480, 242, 146, 66, and 20 kDa, respectively. A buffer containing 40 mM Tris-HCl pH 7.5, 50 mM NaCl, 2.5 mM MgCl_2_, and 1 mM DTT was filtered through a 0.22 μm syringe filter (Merck Millex®-GV). 15 μl buffer was added onto the mounted coverslip and the focus was found automatically and stabilized. Single-chain cohesin or STAG1 were added to the droplet to a final concentration of 10 nM, briefly mixed by pipetting and acquired immediately. Experiments were performed at room temperature (22°C). When single-chain cohesin was measured together with STAG1, single-chain cohesin and STAG1 were mixed in an equimolar ratio at their respective stock concentrations on ice without prolonged incubation and then immediately added to the droplet to yield a final concentration of 10 nM each. Immediately after dilution, a series of 6000 frames was acquired at 1 kHz. All samples were measured three times.

## QUANTIFICATION AND STATISTICAL ANALYSIS

### Quantification and segmentation of kymographs

Quantification of DNA loop size and position was performed as described previously^15^. In brief, the intensity of the loop is normalized to the intensity along the entire DNA molecule and multiplied by the known length of the DNA molecule (48.5 kbp). The loop position was determined as the relative position of the loop from one randomly selected end of the DNA for each molecule but kept constant for all analyses. Half of the loop size at every moment was added to this position. The reported loop position (in kbp) refers thus to the tip of the extruded loop. DNA loop size and position were plotted concomitantly versus time (Figures 1B) and in a two-dimensional plot (Figures S1A and S1B). Segmentation of the kymographs into phases of loop extrusion, loop diffusion, and loop slipping was performed manually. As illustrated in Figure 1C, the time course of the loop size was first segmented into phases of loop growth (loop extrusion), stagnant loop size (loop diffusion), and loop size decrease (loop slipping). Phases in which loops colocalize with the DNA ends (5% of the DNA length at each side) were labelled as ‘stalled’ and excluded from further analysis. Loop extrusion and slipping phases were further divided into segments if the direction changed, as determined by the local slope of the loop position over time. After manual segmentation, the classification of segments into extrusion, diffusion, and slipping phases was done semi-automatically by applying a threshold of was used to distinguish loop diffusion from loop extrusion (loop size increase) and slipping (loop size decrease). The direction of loop extrusion and slipping phases was assigned based on the sign of the loop position derivative with respect to time within each extrusion and slipping phase, respectively.

To determine whether loop extrusion phases proceed asymmetrically or symmetrically, the amount of DNA towards both sides (L_dir1_ and L_dir2_ in Figure 2) were quantified by dividing the relative fluorescence of both DNA segments to the left and right of the loop by the length of the DNA molecule (48.5 kbp), as done previously^2,5,7^.

DNA loop extrusion traces by human cohesin were usually acquired at a time resolution of 200 ms (100 ms exposure time for the DNA and cohesin-JF646 channel, respectively, in the absence of labelled NIPBL-ΔN). To check whether phases of loop extrusion, diffusion, or slipping exist which are too short to be segmented with a time resolution of 200 ms, cohesin-mediated DNA loop extrusion traces were also acquired at a time resolution of 20 ms (only the DNA channel was imaged). This data set was then segmented and classified using the time resolution of 20 ms. Additionally, the same data set was subsampled by averaging 5 subsequent frames of the DNA channel (thus effectively increasing the exposure time to 100 ms) and omitting the following 5 frames (thus mimicking the imaging of cohesin-JF646 for another 100 ms). The subsampled data set was segmented and classified independently. The segmentation results of both data sets were quantitatively compared to check for short phases in the 20 ms data set which were missed in the subsampled data set (Figure S1C-S1F). 76 ± 17 % (probability and 95% binomial confidence interval) of all traces were segmented into an equal number of phases (Figure S1C; Jaccard index of 0.80 [N_union_ = 184, N_intersection_ = 148]) and the change points were set, on average, at the same time (mean time difference 0.14 s; Figure S1D) with low variation between individual change points (SD = 1 s; Figure S1D). The fraction of uni-vs. bidirectional traces was identical (Figure S1E) and there was no significant change between the duration of loop extrusion, diffusion, and slipping phases (Figure S1F). We conclude that a time resolution of 200 ms is sufficient to segment traces of DNA loop extrusion into phases of loop extrusion, diffusion, and slipping and that no significantly shorter phases exist.

### Quantification of DNA loop extrusion, loop diffusion, and loop slipping kinetics

The following metrics were calculated which characterize DNA loop extrusion traces of all eukaryotic SMC complexes: loop extrusion rate, loop slipping rate, diffusion constant, loop lifetime, the fraction of unidirectional traces, frequency of direction change, and the fraction of diffusing and slipping loops.

The loop extrusion (Figure S3H) and loop slipping rate (Figure S4I) were determined *via* a linear fit to the loop size across the loop extrusion or loop slipping phase, respectively. To quantify the diffusion constant of loops during diffusion phases (Figure S4H), the mean squared displacement (MSD) of the loop during the diffusion phase was calculated. The diffusion constant was subsequently determined *via* linear regression of the form MSD(τ) = Dτ + o (diffusion constant D, time lag τ, and offset o due to finite localization uncertainty) to the first 10% of the available time lags for sufficient data points of the fit and to avoid flattening of the MSD curve when the loop reaches the DNA ends.

The loop lifetime (Figures 3C and 4C) was determined as the interval from the beginning of the first loop extrusion phase until the complete dissociation of the loop. In cases in which acquisition was stopped before loop dissociation, the loop lifetime was computed from the beginning of loop extrusion until the end of acquisition.

A trace was labelled as extruding toward one direction only (unidirectional; Figures 1E, 3A, and 4A) if a trace showed at least two extrusion phases in only one direction and no extrusion phase in the other direction. A trace was labelled as extruding bidirectionally if the trace exhibited at least one extrusion phase in each direction. Traces which exhibited only one extrusion phase were not considered because it is unclear whether the SMC complex could only undergo a single phase of extrusion despite being a bidirectional extruder or whether the SMC complex was indeed a unidirectional extruder. The average frequency of direction changes was computed for each loop by dividing the number of direction switches per trace by the loop lifetime (Figures 3B and 4B) for traces which showed at least one direction switch.

The fraction of traces which showed loop diffusion (Figure S3C) and loop slipping (Figure S3D) was computed as the fraction of traces with at least one diffusion or slipping phase, respectively.

### Bleaching and annotation of NIPBL-ΔN presence

The intensity of cohesin-JF646 (in bleaching experiments; Figure S3E-S3G) and NIPBL-ΔN-A550/JF646 was computed as the average intensity in the respective channel at the loop position over a window of 5 pixels around the loop centroid position. The number of bleaching steps of cohesin-JF646 was determined by plotting the cohesin-JF646 intensity at the loop over time and counting the number of step-wise decreases to the background level (Figure S3E). Only cohesin complexes associated with a DNA loop were considered, which allows an assignment of the number of cohesin-JF646 bleaching steps to traces showing uni-(Figure S3F) or bidirectional extrusion (Figure S3G). The appearance and disappearance of NIPBL-ΔN-A550/JF646 (Figures 3H-3M, 4F-4I, and S4B-S4I) was assessed analogously.

### Testing the Markov property

Let us denote the state *k* of cohesin as being in an extrusion, diffusion, or slipping phase. A finite, first-order, discrete Markov chain is a stochastic process in which the probability to transition from state *k* to state *l* depends only on the current state (state *k*) but on states at previous points in time^70^: 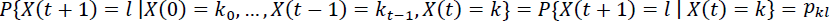. The Markov chain is thus determined by the transition matrix Π which summarizes all 3x3 possible transitions (here we consider three possible states):

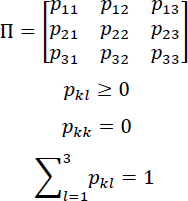

The transition probability *p_ki_* is computed as the number of transitions from state *k* to state *l*, *N*, divided by the number of transitions from state *k* to any state, i.e. 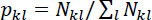.

Alternatively, the stochastic process might have memory, in which case the transition probability from state *k* (at time t) to state *l* (at time t+1) depends also on the previous state *j* (at time t-1), i.e. 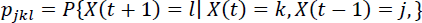. A three-state transition matrix is constructed analogously to the two-state transition matrix Π. The chi-square statistic is subsequently used to test whether the Markov chain is a first order (memory-less, the null hypothesis H_0_) or second order (with memory, H_1_) process^71^. The chi-square statistic for the null hypothesis of state *k* (in other words: the hypothesis that the transition from the current state *k* to the future state *l* is independent of the past state *j*) is:

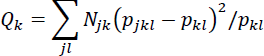

Where 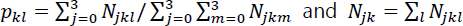. If H_0_ is true, *Q_k_* has a chi-square distribution with (3-1)^2^ degrees of freedom. Similarly, the joint hypothesis that the transition from all states is memory-less, i.e. that 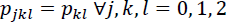, can be computed via the sum 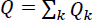 which has a chi-square distribution with 3(2-1)^2^ degrees of freedom. The null hypothesis H_0_ is rejected if the test result *P*(*X* > *Q_k_*) is significant at the significance level of α = 0.05. The p-values for the three states are all > 0.05 across all conditions (Figure S3A and S3B), thus cohesin switches from one state to another without memory of the past state.

### Protein complex structure prediction using AlphaFold2

AlphaFold2 predictions were generated using ColabFold, running on Google Colab^72^ using the following parameters: msa_mode=mmseqs2_uniref_env; pair_mode=unpaired_paired; num_models=1 or 5 (predictions of large complexes quickly run out of GPU memory in which case only 1 model could be predicted); num_recycles=3; recycle_early_stop_tolerance=auto; max_msa=auto; num_seeds=1; use_dropout=True. The sequences used to generate the predictions are listed in Supplementary Note 1. Protein structures were rendered using UCSF ChimeraX (Figures 5A-5F, ref. ^73^).

## ACKNOWLEDGEMENTS

We thank Brian Analikwu and Richard Janissen for discussions and comments on the manuscript and Alejandro Martin Gonzalez, Allard Katan, and Miloš Tišma for discussions. We thank Mathias Madalinski for purifying Atto550- and JF646-HaloTag Ligands. Research in the laboratory of C.D. was supported by ERC Advanced Grant 883684 (DNA looping) and the BaSyC program. Research in the laboratory of J.-M.P. was supported by Boehringer Ingelheim, the Austrian Research Promotion Agency (Headquarter grant FFG-FO999902549), the European Research Council under the European Union’s Horizon 2020 research and innovation program GA No 101020558, the Human Frontier Science Program (grant RGP0057/2018) and the Vienna Science and Technology Fund (grant LS19-029). J.-M.P. is also an adjunct professor at the Medical University of Vienna. Research in the laboratory of S.G. was supported by ERC Consolidator Grant 724482.

## AUTHOR CONTRIBUTIONS

R.B., J.v.d.T., and C.D. designed experiments; R.B. conducted experiments, analysed data, and performed AlphaFold2 predictions; I.D. purified human cohesin, NIPBL-Mau2, STAG1, fluorescently labelled NIPBL-ΔN, and established dual-color NIPBL-ΔN imaging; J.v.d.T. made the DNA constructs, M.T. purified yeast SMC5/6; R.B. and C.D. wrote the manuscript with input from all authors; S.G., J.-M.P., and C.D. acquired funding and supervised the work.

## RESOURCE AVAILABILITY

Further information and requests for resources and reagents should be directed to the lead contact, Cees Dekker.

### Data and code availability

All data reported in this paper will be shared by the lead contact upon request. Custom code will be made publicly available upon publication.

### Data availability

All data of this study are available upon request from the corresponding author Cees Dekker.

## REFERENCES

1. Kim, E., Barth, R., and Dekker, C. (2023). Looping the Genome with SMC Complexes. Annu. Rev. Biochem. 92, 15–41. 10.1146/annurev-biochem-032620-110506.

2. Ganji, M., Shaltiel, I.A., Bisht, S., Kim, E., Kalichava, A., Haering, C.H., and Dekker, C. (2018). Real-time imaging of DNA loop extrusion by condensin. Science 360, 102–105. 10.1126/science.aar7831.

3. Davidson, I.F., Bauer, B., Goetz, D., Tang, W., Wutz, G., and Peters, J.M. (2019). DNA loop extrusion by human cohesin. Science 366, 1338–1345. 10.1126/science.aaz3418.

4. Kim, Y., Shi, Z., Zhang, H., Finkelstein, I.J., and Yu, H. (2019). Human cohesin compacts DNA by loop extrusion. Science 366, 1345–1349. 10.1126/science.aaz4475.

5. Golfier, S., Quail, T., Kimura, H., and Brugués, J. (2020). Cohesin and condensin extrude DNA loops in a cell-cycle dependent manner. eLife 9, 1–34. 10.7554/eLife.53885.

6. Higashi, T.L., Pobegalov, G., Tang, M., Molodtsov, M.I., and Uhlmann, F. (2021). A brownian ratchet model for dna loop extrusion by the cohesin complex. eLife 10, 2021.02.14.431132. 10.7554/eLife.67530.

7. Pradhan, B., Kanno, T., Umeda Igarashi, M., Loke, M.S., Baaske, M.D., Wong, J.S.K., Jeppsson, K., Björkegren, C., and Kim, E. (2023). The Smc5/6 complex is a DNA loop-extruding motor. Nature 616, 843–848. 10.1038/s41586-023-05963-3.

8. Tang, M., Pobegalov, G., Tanizawa, H., Chen, Z.A., Rappsilber, J., Molodtsov, M., Noma, K., and Uhlmann, F. (2023). Establishment of dsDNA-dsDNA interactions by the condensin complex. Mol. Cell, S1097276523007463. 10.1016/j.molcel.2023.09.019.

9. Janissen, R., Barth, R., Davidson, I.F., Taschner, M., Gruber, S., Peters, J.M., and Dekker, C. (2023). All eukaryotic SMC proteins induce a twist of -0.6 at each DNA-loop-extrusion step. bioRxiv.

10. Kong, M., Cutts, E.E., Pan, D., Beuron, F., Kaliyappan, T., Xue, C., Morris, E.P., Musacchio, A., Vannini, A., and Greene, E.C. (2020). Human Condensin I and II Drive Extensive ATP-Dependent Compaction of Nucleosome-Bound DNA. Mol. Cell 79, 99–114.e9. 10.1016/j.molcel.2020.04.026.

11. Ryu, J.-K., Rah, S.-H., Janissen, R., Kerssemakers, J.W.J., Bonato, A., Michieletto, D., and Dekker, C. (2022). Condensin extrudes DNA loops in steps up to hundreds of base pairs that are generated by ATP binding events. Nucleic Acids Res. 50, 820–832. 10.1093/nar/gkab1268.

12. Goloborodko, A., Imakaev, M.V., Marko, J.F., and Mirny, L. (2016). Compaction and segregation of sister chromatids via active loop extrusion. eLife 5. 10.7554/eLife.14864.

13. Gibcus, J.H., Samejima, K., Goloborodko, A., Samejima, I., Naumova, N., Nuebler, J., Kanemaki, M.T., Xie, L., Paulson, J.R., Earnshaw, W.C., et al. (2018). A pathway for mitotic chromosome formation. Science 359. 10.1126/science.aao6135.

14. de Wit, E., Vos, E.S.M., Holwerda, S.J.B., Valdes-Quezada, C., Verstegen, M.J.A.M., Teunissen, H., Splinter, E., Wijchers, P.J., Krijger, P.H.L., and de Laat, W. (2015). CTCF Binding Polarity Determines Chromatin Looping. Mol. Cell 60, 676–684. 10.1016/j.molcel.2015.09.023.

15. Davidson, I.F., Barth, R., Zaczek, M., Van Der Torre, J., Tang, W., Nagasaka, K., Janissen, R., Kerssemakers, J., Wutz, G., Dekker, C., et al. (2023). CTCF is a DNA-tension-dependent barrier to cohesin-mediated loop extrusion. Nature 616, 822–827. 10.1038/s41586-023-05961-5.

16. Guo, Y., Xu, Q., Canzio, D., Shou, J., Li, J., Gorkin, D.U., Jung, I., Wu, H., Zhai, Y., Tang, Y., et al. (2015). CRISPR Inversion of CTCF Sites Alters Genome Topology and Enhancer/Promoter Function. Cell 162, 900–910. 10.1016/j.cell.2015.07.038.

17. Li, Y., Haarhuis, J.H.I., Sedeño Cacciatore, Á., Oldenkamp, R., van Ruiten, M.S., Willems, L., Teunissen, H., Muir, K.W., de Wit, E., Rowland, B.D., et al. (2020). The structural basis for cohesin– CTCF-anchored loops. Nature 578, 472–476. 10.1038/s41586-019-1910-z.

18. Nora, E.P., Lajoie, B.R., Schulz, E.G., Giorgetti, L., Okamoto, I., Servant, N., Piolot, T., Van Berkum, N.L., Meisig, J., Sedat, J., et al. (2012). Spatial partitioning of the regulatory landscape of the X-inactivation centre. Nature 485, 381–385. 10.1038/nature11049.

19. Nora, E.P., Goloborodko, A., Valton, A.L., Gibcus, J.H., Uebersohn, A., Abdennur, N., Dekker, J., Mirny, L.A., and Bruneau, B.G. (2017). Targeted Degradation of CTCF Decouples Local Insulation of Chromosome Domains from Genomic Compartmentalization. Cell 169, 930–944.e22. 10.1016/j.cell.2017.05.004.

20. Rao, S.S.P., Huntley, M.H., Durand, N.C., Stamenova, E.K., Bochkov, I.D., Robinson, J.T., Sanborn, A.L., Machol, I., Omer, A.D., Lander, E.S., et al. (2014). A 3D map of the human genome at kilobase resolution reveals principles of chromatin looping. Cell 159, 1665–1680. 10.1016/j.cell.2014.11.021.

21. Rao, S.S.P., Huang, S.C., Glenn St Hilaire, B., Engreitz, J.M., Perez, E.M., Kieffer-Kwon, K.R., Sanborn, A.L., Johnstone, S.E., Bascom, G.D., Bochkov, I.D., et al. (2017). Cohesin Loss Eliminates All Loop Domains. Cell 171, 305–320.e24. 10.1016/j.cell.2017.09.026.

22. Vietri Rudan, M., Barrington, C., Henderson, S., Ernst, C., Odom, D.T., Tanay, A., and Hadjur, S. (2015). Comparative Hi-C Reveals that CTCF Underlies Evolution of Chromosomal Domain Architecture. Cell Rep. 10, 1297–1309. 10.1016/j.celrep.2015.02.004.

23. Hansen, A.S., Cattoglio, C., Darzacq, X., and Tjian, R. (2018). Recent evidence that TADs and chromatin loops are dynamic structures. Nucleus 9, 20–32. 10.1080/19491034.2017.1389365.

24. Gabriele, M., Brandão, H.B., Grosse-Holz, S., Jha, A., Dailey, G.M., Cattoglio, C., Hsieh, T.H.S., Mirny, L., Zechner, C., and Hansen, A.S. (2022). Dynamics of CTCF- and cohesin-mediated chromatin looping revealed by live-cell imaging. Science 376, 476–501. 10.1126/SCIENCE.ABN6583/SUPPL_FILE/SCIENCE.ABN6583_MDAR_REPRODUCIBILITY_CHECKLIST.PDF.

25. Mach, P., Kos, P.I., Zhan, Y., Cramard, J., Gaudin, S., Tünnermann, J., Marchi, E., Eglinger, J., Zuin, J., Kryzhanovska, M., et al. (2022). Cohesin and CTCF control the dynamics of chromosome folding. Nat. Genet. 54, 1907–1918. 10.1038/s41588-022-01232-7.

26. Horsfield, J.A. (2022). Full circle: a brief history of cohesin and the regulation of gene expression. FEBS J., febs.16362. 10.1111/febs.16362.

27. Arnould, C., Rocher, V., Finoux, A.L., Clouaire, T., Li, K., Zhou, F., Caron, P., Mangeot, P.E., Ricci, E.P., Mourad, R., et al. (2021). Loop extrusion as a mechanism for formation of DNA damage repair foci. Nature 590, 660–665. 10.1038/s41586-021-03193-z.

28. Litwin, I., Pilarczyk, E., and Wysocki, R. (2018). The Emerging Role of Cohesin in the DNA Damage Response. Genes 9, 581. 10.3390/genes9120581.

29. Phipps, J., Nasim, A., and Miller, D.R. (1985). Recovery, Repair, and Mutagenesis in Schizosaccharomyces pombe. In Advances in Genetics (Elsevier), pp. 1–72. 10.1016/S0065-2660(08)60511-8.

30. Aragón, L. (2018). The Smc5/6 Complex: New and Old Functions of the Enigmatic Long-Distance Relative. Annu. Rev. Genet. 52, 89–107. 10.1146/annurev-genet-120417-031353.

31. McDonald, W.H., Pavlova, Y., Yates, J.R., and Boddy, M.N. (2003). Novel Essential DNA Repair Proteins Nse1 and Nse2 Are Subunits of the Fission Yeast Smc5-Smc6 Complex. J. Biol. Chem. 278, 45460–45467. 10.1074/jbc.M308828200.

32. Bürmann, F., and Löwe, J. (2023). Structural biology of SMC complexes across the tree of life. Curr. Opin. Struct. Biol. 80, 102598. 10.1016/j.sbi.2023.102598.

33. Datta, S., Lecomte, L., and Haering, C.H. (2020). Structural insights into DNA loop extrusion by SMC protein complexes. Curr. Opin. Struct. Biol. 65, 102–109. 10.1016/j.sbi.2020.06.009.

34. Oldenkamp, R., and Rowland, B.D. (2022). A walk through the SMC cycle: From catching DNAs to shaping the genome. Mol. Cell 82, 1616–1630. 10.1016/j.molcel.2022.04.006.

35. Shaltiel, I.A., Datta, S., Lecomte, L., Hassler, M., Kschonsak, M., Bravo, S., Stober, C., Ormanns, J., Eustermann, S., and Haering, C.H. (2022). A hold-and-feed mechanism drives directional DNA loop extrusion by condensin. Science 376, 1087–1094. 10.1126/science.abm4012.

36. Dekker, C., Haering, C.H., Peters, J.-M., and Rowland, B.D. (2023). How do molecular motors fold the genome? Science 382, 646–648. 10.1126/science.adi8308.

37. Davidson, I.F., Barth, R., Zaczek, M., Van Der Torre, J., Tang, W., Nagasaka, K., Janissen, R., Kerssemakers, J., Wutz, G., Dekker, C., et al. (2023). CTCF is a DNA-tension-dependent barrier to cohesin-mediated loop extrusion. Nature 616, 822–827. 10.1038/s41586-023-05961-5.

38. Taschner, M., and Gruber, S. (2023). DNA segment capture by Smc5/6 holocomplexes. Nat. Struct. Mol. Biol. 30, 619–628. 10.1038/s41594-023-00956-2.

39. Kschonsak, M., Merkel, F., Bisht, S., Metz, J., Rybin, V., Hassler, M., and Haering, C.H. (2017). Structural Basis for a Safety-Belt Mechanism That Anchors Condensin to Chromosomes. Cell 171, 588–600.e24. 10.1016/j.cell.2017.09.008.

40. Rhodes, J., Mazza, D., Nasmyth, K., and Uphoff, S. (2017). Scc2/Nipbl hops between chromosomal cohesin rings after loading. eLife 6, e30000. 10.7554/eLife.30000.

41. Bauer, B.W., Davidson, I.F., Canena, D., Wutz, G., Tang, W., Litos, G., Horn, S., Hinterdorfer, P., and Peters, J.M. (2021). Cohesin mediates DNA loop extrusion by a “swing and clamp” mechanism. Cell 184, 5448–5464. 10.1016/j.cell.2021.09.016.

42. Shi, Z., Gao, H., Bai, X.C., and Yu, H. (2020). Cryo-EM structure of the human cohesin-NIPBL-DNA complex. Science 368, 1454–1459. 10.1126/science.abb0981.

43. Collier, J.E., Lee, B.G., Roig, M.B., Yatskevich, S., Petela, N.J., Metson, J., Voulgaris, M., Llamazares, A.G., Löwe, J., and Nasmyth, K.A. (2020). Transport of DNA within cohesin involves clamping on top of engaged heads by SCC2 and entrapment within the ring by SCC3. eLife 9, 1–36. 10.7554/ELIFE.59560.

44. Takaki, R., Dey, A., Shi, G., and Thirumalai, D. (2021). Theory and simulations of condensin mediated loop extrusion in DNA. Nat. Commun. 12, 5865. 10.1038/s41467-021-26167-1.

45. Davidson, I.F., and Peters, J.M. (2021). Genome folding through loop extrusion by SMC complexes. Nat. Rev. Mol. Cell Biol. 22, 445–464. 10.1038/s41580-021-00349-7.

46. Lee, B.G., Merkel, F., Allegretti, M., Hassler, M., Cawood, C., Lecomte, L., O’Reilly, F.J., Sinn, L.R., Gutierrez-Escribano, P., Kschonsak, M., et al. (2020). Cryo-EM structures of holo condensin reveal a subunit flip-flop mechanism. Nat. Struct. Mol. Biol. 27, 743–751. 10.1038/s41594-020-0457-x.

47. Jumper, J., Evans, R., Pritzel, A., Green, T., Figurnov, M., Ronneberger, O., Tunyasuvunakool, K., Bates, R., Žídek, A., Potapenko, A., et al. (2021). Highly accurate protein structure prediction with AlphaFold. Nature 596, 583–589. 10.1038/s41586-021-03819-2.

48. Murayama, Y., and Uhlmann, F. (2014). Biochemical reconstitution of topological DNA binding by the cohesin ring. Nature 505, 367–371. 10.1038/nature12867.

49. Nasmyth, K.A., Lee, B.-G., Roig, M.B., and Löwe, J. (2023). What AlphaFold tells us about cohesin’s retention on and release from chromosomes (Biochemistry) 10.1101/2023.04.14.536858.

50. Holzmann, J., Politi, A.Z., Nagasaka, K., Hantsche-Grininger, M., Walther, N., Koch, B., Fuchs, J., Dürnberger, G., Tang, W., Ladurner, R., et al. (2019). Absolute quantification of cohesin, CTCF and their regulators in human cells. eLife 8, e46269. 10.7554/eLife.46269.

51. Huang, Q., Szklarczyk, D., Wang, M., Simonovic, M., and Von Mering, C. (2023). PaxDb 5.0: Curated Protein Quantification Data Suggests Adaptive Proteome Changes in Yeasts. Mol. Cell. Proteomics 22, 100640. 10.1016/j.mcpro.2023.100640.

52. Ho, B., Baryshnikova, A., and Brown, G.W. (2018). Unification of Protein Abundance Datasets Yields a Quantitative Saccharomyces cerevisiae Proteome. Cell Syst. 6, 192–205.e3. 10.1016/j.cels.2017.12.004.

53. Cattoglio, C., Pustova, I., Walther, N., Ho, J.J., Hantsche-Grininger, M., Inouye, C.J., Hossain, M.J., Dailey, G.M., Ellenberg, J., Darzacq, X., et al. (2019). Determining cellular CTCF and cohesin abundances to constrain 3D genome models. eLife 8. 10.7554/elife.40164.

54. Hu, B., Itoh, T., Mishra, A., Katoh, Y., Chan, K.-L., Upcher, W., Godlee, C., Roig, M.B., Shirahige, K., and Nasmyth, K. (2011). ATP Hydrolysis Is Required for Relocating Cohesin from Sites Occupied by Its Scc2/4 Loading Complex. Curr. Biol. 21, 12–24. 10.1016/j.cub.2010.12.004.

55. Fudenberg, G., Abdennur, N., Imakaev, M., Goloborodko, A., and Mirny, L.A. (2017). Emerging Evidence of Chromosome Folding by Loop Extrusion. Cold Spring Harb. Symp. Quant. Biol. 82, 45–55. 10.1101/sqb.2017.82.034710.

56. Banigan, E.J., van den Berg, A.A., Brandão, H.B., Marko, J.F., and Mirny, L.A. (2020). Chromosome organization by one-sided and two-sided loop extrusion. eLife 9, e53558. 10.7554/eLife.53558.

57. Dey, A., Shi, G., Takaki, R., and Thirumalai, D. (2023). Structural changes in chromosomes driven by multiple condensin motors during mitosis. Cell Rep. 42, 112348. 10.1016/j.celrep.2023.112348.

58. Galitsyna, A., Ulianov, S.V., Bykov, N.S., Veil, M., Gao, M., Perevoschikova, K., Gelfand, M., Razin, S.V., Mirny, L., and Onichtchouk, D. (2023). Extrusion fountains are hallmarks of chromosome organization emerging upon zygotic genome activation (Molecular Biology) 10.1101/2023.07.15.549120.

59. Isiaka, B.N., Semple, J.I., Haemmerli, A., Thapliyal, S., Stojanovski, K., Das, M., Gilbert, N., Glauser, D.A., Towbin, B., Jost, D., et al. (2023). Cohesin forms fountains at active enhancers in C. elegans (Genomics) 10.1101/2023.07.14.549011.

60. Liu, N.Q., Magnitov, M., Schijns, M., Van Schaik, T., Van Der Weide, R.H., Teunissen, H., Van Steensel, B., and De Wit, E. (2021). Rapid depletion of CTCF and cohesin proteins reveals dynamic features of chromosome architecture (Molecular Biology) 10.1101/2021.08.27.457977.

61. Guo, Y., Al-Jibury, E., Garcia-Millan, R., Ntagiantas, K., King, J.W.D., Nash, A.J., Galjart, N., Lenhard, B., Rueckert, D., Fisher, A.G., et al. (2022). Chromatin jets define the properties of cohesin-driven in vivo loop extrusion. Mol. Cell 82, 3769–3780.e5. 10.1016/j.molcel.2022.09.003.

62. Kim, E., Kerssemakers, J., Shaltiel, I.A., Haering, C.H., and Dekker, C. (2020). DNA-loop extruding condensin complexes can traverse one another. Nature 579, 438–442. 10.1038/s41586-020-2067-5.

63. Kim, E., Gonzalez, A.M., Pradhan, B., van der Torre, J., and Dekker, C. (2022). Condensin-driven loop extrusion on supercoiled DNA. Nat. Struct. Mol. Biol. 10.1038/s41594-022-00802-x.

64. Pradhan, B., Barth, R., Kim, E., Davidson, I.F., Bauer, B., van Laar, T., Yang, W., Ryu, J.-K., van der Torre, J., Peters, J.-M., et al. (2022). SMC complexes can traverse physical roadblocks bigger than their ring size. Cell Rep. 41, 111491. 10.1016/j.celrep.2022.111491.

65. Park, S.R., Hauver, J., Zhang, Y., Revyakin, A., Coleman, R.A., Tjian, R., Chu, S., and Pertsinidis, A. (2020). A Single-Molecule Surface-Based Platform to Detect the Assembly and Function of the Human RNA Polymerase II Transcription Machinery. Structure 28, 1337–1343.e4. 10.1016/j.str.2020.07.009.

66. Jeppsson, K., Pradhan, B., Sutani, T., Sakata, T., Igarashi, M.U., Berta, D.G., Kanno, T., Nakato, R., Shirahige, K., Kim, E., et al. (2023). Loop-extruding Smc5/6 organizes transcription-induced positive DNA supercoils (Molecular Biology) 10.1101/2023.06.20.545053.

67. Martínez-García, B., Dyson, S., Segura, J., Ayats, A., Cutts, E.E., Gutierrez-Escribano, P., Aragón, L., and Roca, J. (2023). Condensin pinches a short negatively supercoiled DNA loop during each round of ATP usage. EMBO J. 42, e111913. 10.15252/embj.2022111913.

68. Kimura, K., and Hirano, T. (1997). ATP-dependent positive supercoiling of DNA by 13S condensin: A biochemical implication for chromosome condensation. Cell 90, 625–634. 10.1016/S0092-8674(00)80524-3.

69. Kapusta, P. (2010). Absolute diffusion coefficients: compilation of reference data for FCS calibration.

70. Kemeny, J.G., and Snell, J.L. (1960). Finite Markov chains (Van Nostrand).

71. Anderson, T.W., and Goodman, L.A. (1957). Statistical Inference about Markov Chains. Ann. Math. Stat. 28, 89–110. 10.1214/aoms/1177707039.

72. Mirdita, M., Schütze, K., Moriwaki, Y., Heo, L., Ovchinnikov, S., and Steinegger, M. (2022). ColabFold: making protein folding accessible to all. Nat. Methods 19, 679–682. 10.1038/s41592-022-01488-1.

73. Goddard, T.D., Huang, C.C., Meng, E.C., Pettersen, E.F., Couch, G.S., Morris, J.H., and Ferrin, T.E. (2018). UCSF ChimeraX: Meeting modern challenges in visualization and analysis. Protein Sci. 27, 14–25. 10.1002/pro.3235.

