## Supplementary Information for "SMC motor proteins extrude DNA asymmetrically and contain a direction switch"

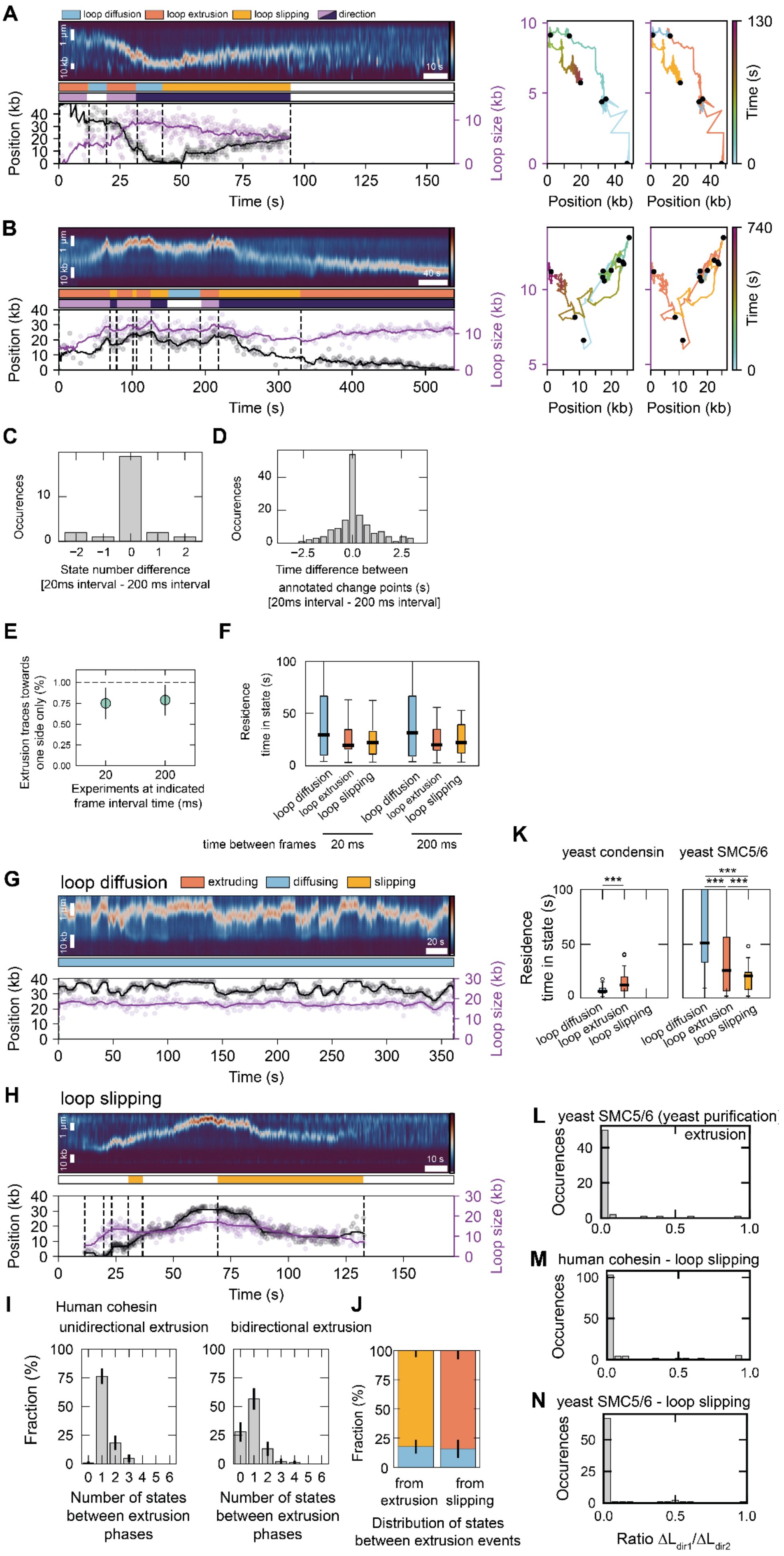
A
loop diffusion
loop extrusion
loop slipping
direction
1 µm
10 kb
10 s
Position (kb)
Loop size (kb)
Time (s)
Position (kb)
Time (s)
130
B
40 s
740
C
Occurences
State number difference
[20ms interval - 200 ms interval
D
Occurences
Time difference between
annotated change points (s)
[20ms interval - 200 ms interval]
E
Extrusion traces towards one side only (%)
Experiments at indicated frame interval time (ms)
F
Residence time in state (s)
loop diffusion
loop extrusion
loop slipping
time between frames
20 ms
200 ms
K
yeast condensin
yeast SMC5/6
Residence time in state (s)
G
loop diffusion
extruding
diffusing
slipping
20 s
H
loop slipping
L
yeast SMC5/6 (yeast purification)
extrusion
Occurences
M
human cohesin - loop slipping
N
yeast SMC5/6 - loop slipping
Ratio ΔL_dir1/ΔL_dir2
I
Human cohesin
unidirectional extrusion
bidirectional extrusion
Fraction (%)
Number of states between extrusion phases
J
from extrusion
from slipping
Distribution of states between extrusion events

**Figure S1: Comparison of DNA loop extrusion trace segmentation between data acquisition with 20 ms and 200 ms time resolution. Loop diffusion and slipping examples, state residence times for yeast SMC5/6 and yeast condensin. Loop slipping is, like extrusion, asymmetric. Related to Figures 1 and 2.**

**(A)** Kymograph of cohesin-mediated DNA loop extrusion (left). Loop position and size are quantified and segmented into loop extrusion, diffusion, and slipping phases. The two panels on the right display a two-dimensional plot of the loop size versus loop position. Black dots demarcate boundaries between diffusion, extrusion, and slipping phases. Diffusion phases move on horizontal lines in this graph, and extrusion and slipping phases are demarcated by diagonal lines.

**(B)** as (A) but for a loop extrusion trace in which extrusion phases in both directions are present as shown in Figure 1B.

**(C)** Difference in the number of phases (segments) between the data set acquired at 20 ms frame interval time and the subsampled one (interval time 200 ms). Negative values indicate that fewer segments were segmented in the 20 ms interval data set than in the 200 ms data set.

**(D)** Time between the annotated change points (time points between segments) in the data set with an interval time of 20 ms and 200 ms between frames. 74% of all change points coincide between the two annotations within 1 s (N = 147 segments). Negative values indicate that the change point in the 20 ms interval data set was annotated prior to the respective change point in the 200 ms data set.

**(E)** Fraction of traces with at least two extrusion events that displayed only unidirectional extrusion (N = 25 traces each). Error bars denote the binomial 95% confidence interval.

**(F)** Duration of diffusion, extrusion, and slipping phases in the annotated data sets acquired at 20 ms frame interval time and the subsampled one (interval time 200 ms). Black horizontal lines are median values, the box extends between the first and third quartile, and the whiskers extend to 1.5*IQR. No statistically significant difference was found between loop diffusion, extrusion, and slipping states between the two data sets. Statistical significance was assessed by a Mann-Whitney test with Bonferroni correction (p-values are 0.9514, 0.6504, and 0.4710 for loop diffusion, extrusion, and slipping states, respectively).

**(G)** Exemplary kymograph of human cohesin showing loop diffusion.

**(H)** Exemplary kymograph of human cohesin showing loop slipping.

**(I)** Duration of diffusion, extrusion, and slipping states for yeast condensin (N = 16, 48, 1, respectively) and yeast SMC5/6 (N = 52, 124, 80, respectively). Black horizontal lines are median values, the box extends between first and third quartile, and whiskers extend to 1.5*IQR. Statistical significance was assessed by a Mann-Whitney test with Bonferroni correction. (***: $p<0.001$).

**(J)** Number of intervening states between extrusion events for traces showing only unidirectional (N = 148) or bidirectional extrusion (N = 108). Bar height denotes the probability of finding the annotated number of steps between extrusion events. Error bars denote the binomial 95% confidence interval.

**(K)** Distribution of states between direction exchange in extrusion (N = 90) and slipping (N = 88) events. Error bars denote the binomial 95% confidence interval.

**(L)** Distribution of the ratio $\Delta L_{dir1}/\Delta L_{dir2}$ for extrusion events of yeast SMC5/6 purified from yeast cells (N = 56 events). Yeast SMC5/6 purified from yeast cells exhibits asymmetric extrusion phases (93 $\pm$ 7 % [mean $\pm$ 95% binomial confidence interval] of $\Delta L_{dir1}/\Delta L_{dir2}$ values below 0.1).

**(M)** Distribution of the ratio $\Delta L_{dir1}/\Delta L_{dir2}$ for slipping events of human cohesin (N = 130 events).

**(N)** Distribution of the ratio $\Delta L_{dir1}/\Delta L_{dir2}$ for slipping events of yeast SMC5/6, purified from *E.coli* (N = 77 events).

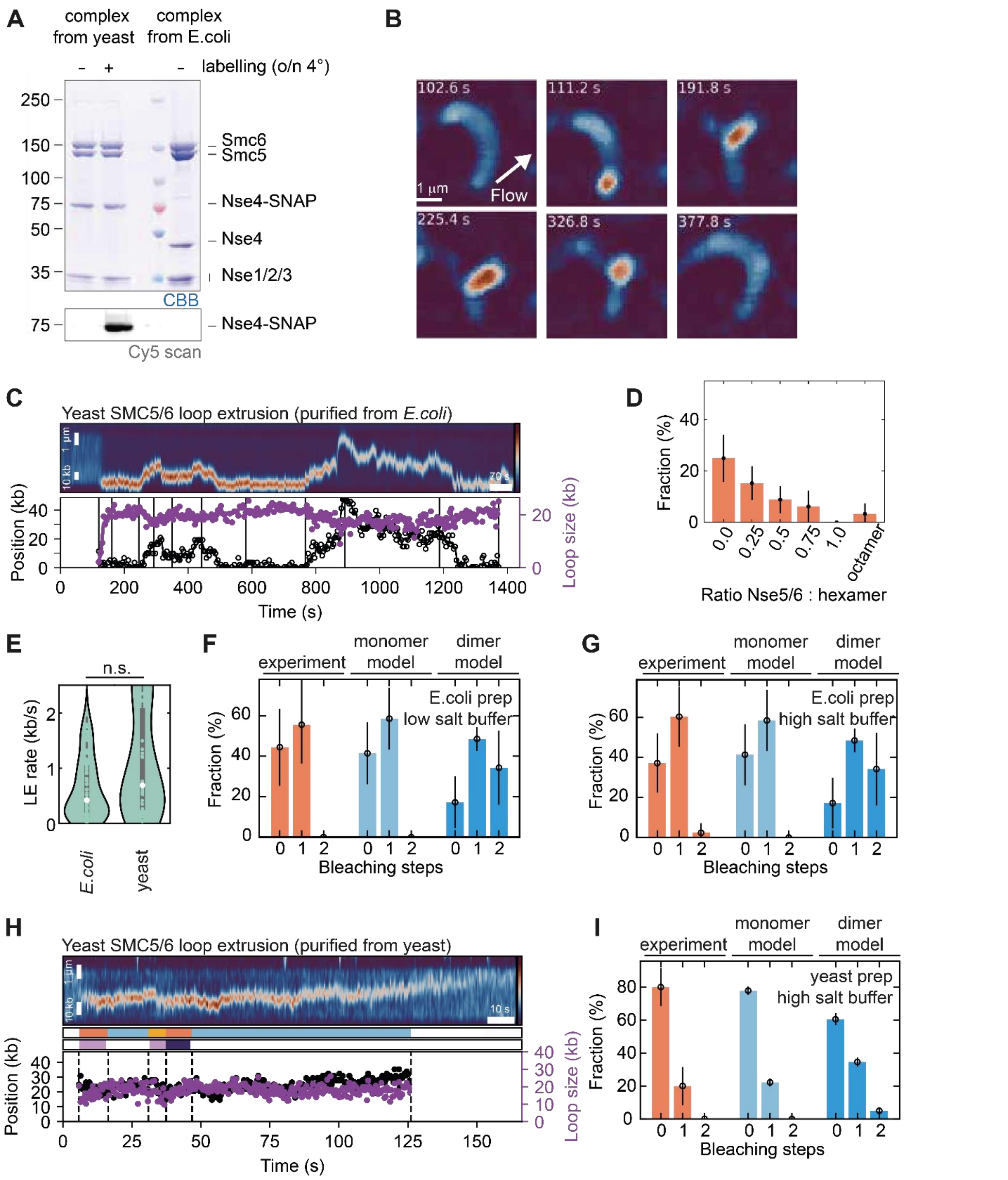


**Figure S2: DNA loop extrusion by single SMC5/6 hexamers. Related to Figure 1.**

(**A**) Detection of SMC5/6 hexamer subunits by SDS–PAGE and Coomassie staining for the unlabeled (1st lane), Nse4-Alexa647-labeled (2nd lane) SMC5/6 complex purified from yeast cells, as well as for the unlabeled SMC5/6 purified from *E.coli* as in [1].

(**B**) Fluorescence imaging snapshots showing DNA loop extrusion by an SMC5/6 hexamer purified from *E.coli* during side flow.

(**C**) Exemplary kymograph of DNA loop extrusion by an SMC5/6 hexamer purified from *E.coli*.

(**D**) Loop extrusion efficiency with varying ratio of Nse5/6 to the SMC5/6 hexamer purified from *E.coli*. No loop extrusion is observed at an equimolar ratio of Nse5/6 and the SMC5/6 hexamer. A co-purification of Nse5/6-bound SMC5/6 hexamer (octamer) shows a small fraction of loop extrusion, potentially due to dissociation of Nse5/6 from the SMC5/6 hexamer that allows loop extrusion. Bar heights denote the probability of finding a DNA with a loop and the error bar denotes the binomial 95% confidence interval (N = 92, 124, 124, 64, 74, 84 DNA molecules from left to right).
(**E**) DNA loop extrusion rate of the SMC5/6 hexamer, purified from *E.coli* (ref. [1]) and from yeast. The white dot marks the median value (0.4 $\pm$ 0.5, median $\pm$ SD, N = 83 for *E.coli*; 0.7 $\pm$ 0.9, median $\pm$ SD, N = 30 for *E.coli*), the box extends to the quartiles, and the whiskers denote the interquartile range. Statistical significance was assessed by a two-sided Kolmogorov-Smirnoff test (p = 0.087).
(**F**) Bleaching step distribution of JF646-labeled SMC5/6 hexamers purified from *E.coli* in 50 mM NaCl, 2.5 mM $MgCl_2$ with a labeling efficiency of 58 ± 11% (N = 27 for the experiment; N = 1,000 simulated monomer/dimer complexes). The monomer model fits the experimental data while the dimer model does not. Error bars denote the 95% binomial confidence interval for experimental data and the range of the predicted monomer/dimer bleaching step distributions for a labeling efficiency of 47% (lower end) and 69% (upper end).
(**G**) As for (F) but in a 100 mM NaCl, 7.5 mM $MgCl_2$ buffer, which is the buffer in which dimers of SMC5/6 hexamers were reported to extrude by Pradhan *et al.* (ref. [2]).
(**H**) Exemplary kymograph of DNA loop extrusion by an SMC5/6 hexamer purified from yeast.
(**I**) As for (G) but for SMC5/6 purified from yeast with a labeling efficiency of 22 ± 2% (N = 50 for the experiment; N = 1,000 simulated monomer/dimer complexes) in a 100 mM NaCl, 7.5 mM $MgCl_2$ buffer, in which dimers of SMC5/6 hexamers were reported to extrude by Pradhan *et al.* (ref. [2]). Error bars denote the 95% binomial confidence interval for experimental data and the range of the predicted monomer/dimer bleaching step distributions for a labeling efficiency of 20% (lower end) and 24% (upper end).

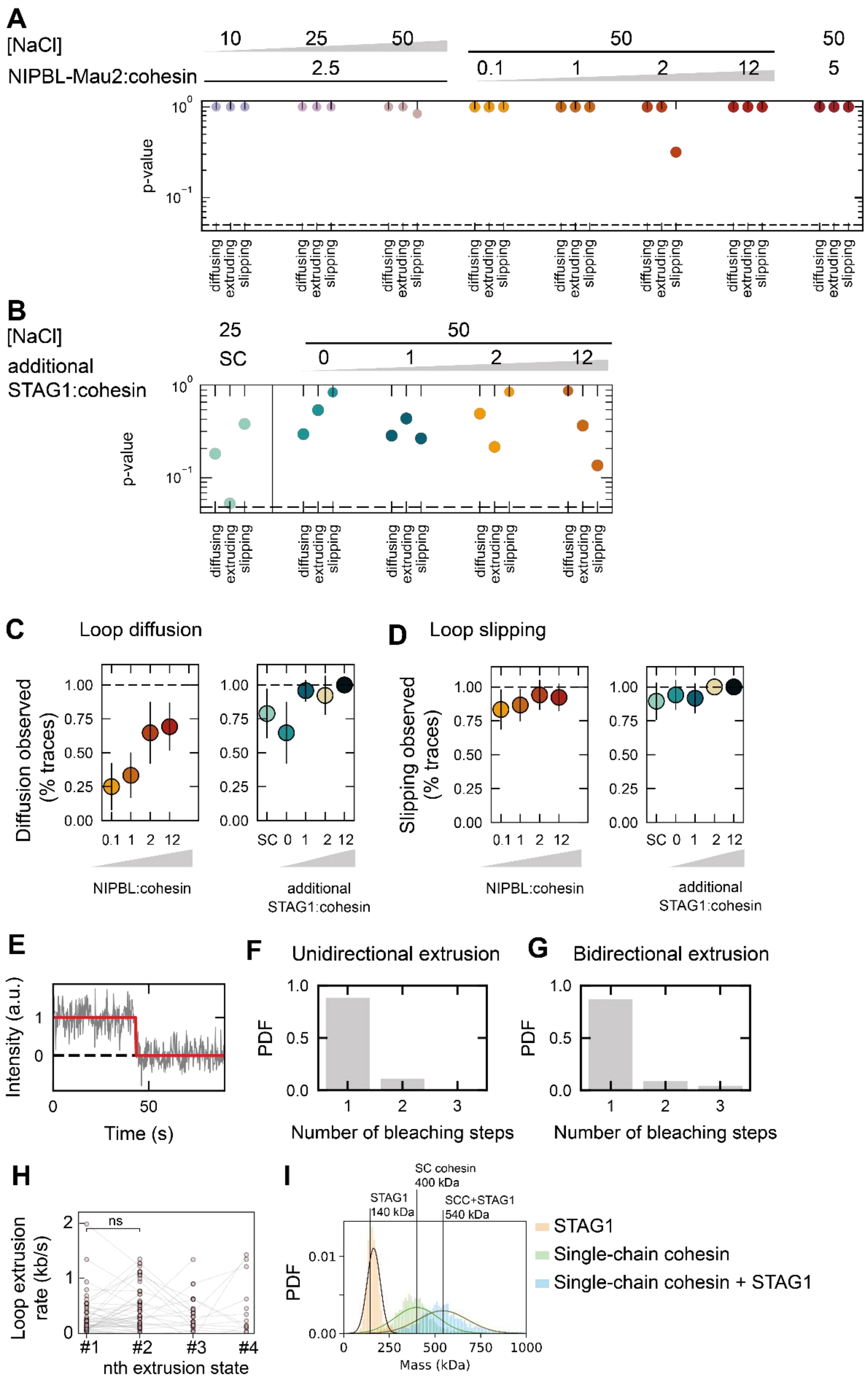
A
[NaCl]
NIPBL-Mau2:cohesin
10
25
50
2.5
50
0.1
1
2
12
50
5
p-value
diffusing
extruding
slipping
B
[NaCl]
additional STAG1:cohesin
25
SC
50
0
1
2
12
p-value
C
Loop diffusion
Diffusion observed (% traces)
NIPBL:cohesin
additional STAG1:cohesin
D
Loop slipping
Slipping observed (% traces)
NIPBL:cohesin
additional STAG1:cohesin
E
Intensity (a.u.)
Time (s)
F
Unidirectional extrusion
PDF
Number of bleaching steps
G
Bidirectional extrusion
PDF
Number of bleaching steps
H
Loop extrusion rate (kb/s)
ns
nth extrusion state
#1
#2
#3
#4
I
STAG1 140 kDa
SC cohesin 400 kDa
SCC+STAG1 540 kDa
PDF
Mass (kDa)
STAG1
Single-chain cohesin
Single-chain cohesin + STAG1

**Figure S3: The sequence of loop diffusion, extrusion, and slipping events is a Markov chain and the excess of NIPBL-Mau2 and STAG1 over cohesin modulate the occurrence of diffusion and slipping phases. Bidirectional extrusion is the result of monomeric cohesin complexes. Related to Figures 1 and 3.**

**(A)** p-value of the Markov property test for transitions starting from a loop diffusion, extrusion, and slipping state for all tested NaCl concentrations and NIPBL-Mau2:cohesin ratios. A value below 0.05 (dashed lin(E) indicates the violation of the Markov property and would suggest a system with memory.

**(B)** As for (A) but for increasing ratios of STAG1:cohesin at constant NIPBL-Mau2:cohesin ratio of 2.5 and 50 mM NaCl. The p-value of the single-chain cohesin (S(C) extrusion data point is 0.054.

**(C)** Fraction of diffusing cohesin complexes for varying NaCl concentrations (NIPBL-Mau2:cohesin = 2.5, no additional STAG1; N = 9, 20, 13 from left to right), for varying ratios of NIPBL-Mau2:cohesin (50 mM NaCl, no additional STAG1; N = 24, 30, 17, 26from left to right), and for single-chain cohesin without STAG1 (NIPBL-Mau2:cohesin = 2.5, 25 mM NaCl) as well as for increasing excess of STAG1 (NIPBL-Mau2:cohesin = 2.5, 50 mM NaCl; N= 19, 17, 24, 13, 30). Dots denote the fraction, error bars denote the binomial 95% confidence interval.

**(D)** As for (C) but for loop slipping. Number of data points as in (C).

**(E)** Exemplary bleaching trace showing a single bleaching step.

**(F)** Number of bleaching steps of human cohesin-JF646 for complexes which extrude unidirectionally (N = 128).

**(G)** Number of bleaching steps of human cohesin-JF646 for complexes which extrude bidirectionally (N = 23).

**(H)** Loop extrusion rate for the $n^{th}$ extrusion state within single kymographs.

**(I)** Mass distribution of STAG1 (orange, N = 735), single-chain cohesin (green, N = 3760), and single-chain cohesin with STAG1 in a 1:1 ratio (blue, N = 2259). The molecular weights of the single and combined complexes are indicated by solid lines. Single-chain cohesin is STAG1-unbound as addition of STAG1 in a 1:1 ratio results in a complex of mass equal to the sum of masses of single-chain cohesin and STAG1.

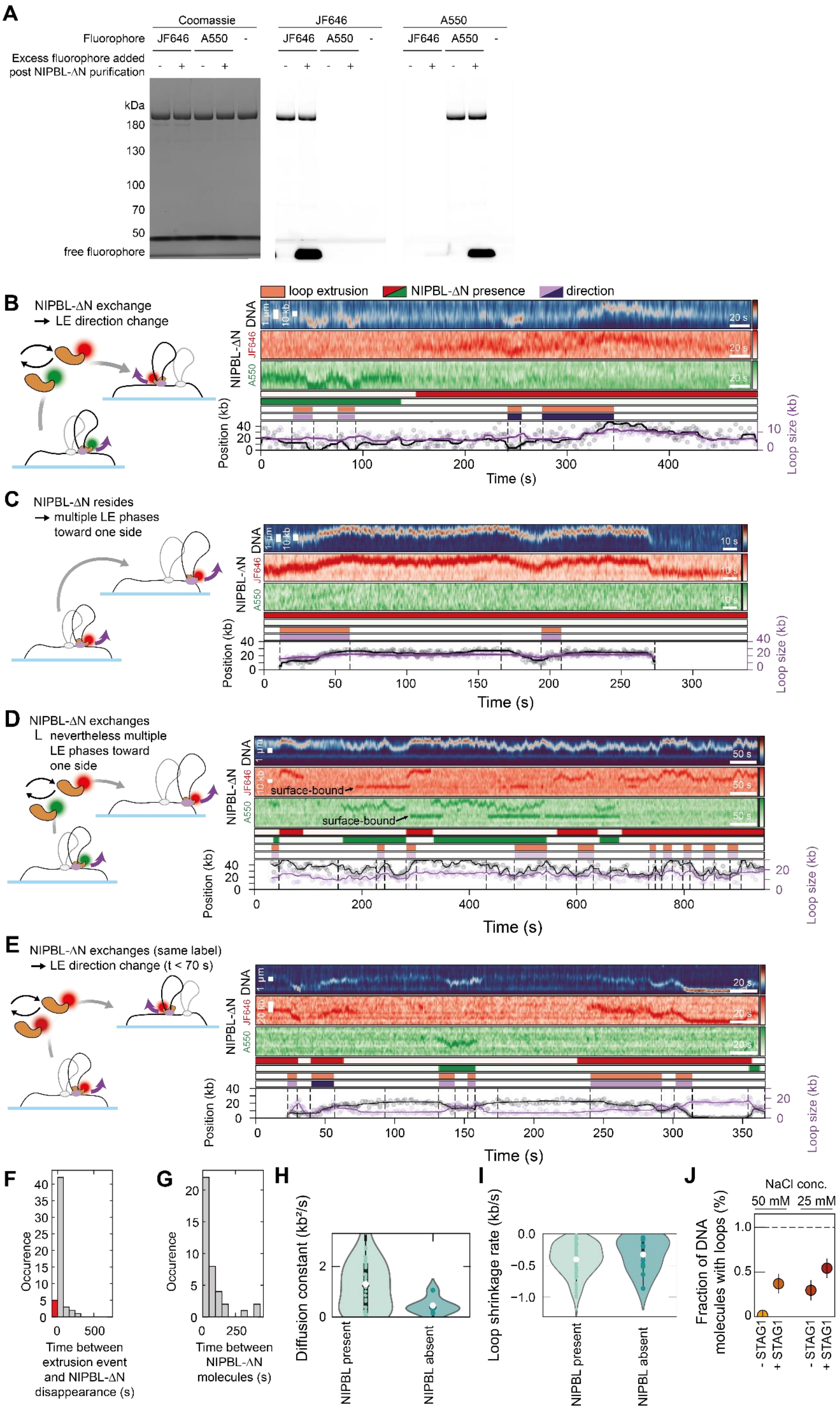
A
Coomassie
JF646
A550
Fluorophore
JF646 A550 -
Excess fluorophore added post NIPBL-ΔN purification
kDa
180
130
100
70
50
free fluorophore
B
NIPBL-ΔN exchange → LE direction change
loop extrusion
NIPBL-ΔN presence
direction
DNA
NIPBL-ΔN
Position (kb)
Loop size (kb)
Time (s)
C
NIPBL-ΔN resides → multiple LE phases toward one side
D
NIPBL-ΔN exchanges nevertheless multiple LE phases toward one side
surface-bound
E
NIPBL-ΔN exchanges (same label) → LE direction change (t < 70 s)
F
Occurence
Time between extrusion event and NIPBL-ΔN disappearance (s)
G
Occurence
Time between NIPBL-ΔN molecules (s)
H
Diffusion constant (kb²/s)
NIPBL present
NIPBL absent
I
Loop shrinkage rate (kb/s)
NIPBL present
NIPBL absent
J
NaCl conc.
50 mM
25 mM
Fraction of DNA molecules with loops (%)
- STAG1
+ STAG1

**Figure S4: Additional loop extrusion traces with differentially labeled NIPBL-ΔN. Related to Figures 3 and 4.**

**(A)** Purity and labeling of NIPBL-ΔN with JF646 and ATTO550, respectively. Molecular weights are indicated in kilo Daltons (kDa). An excess of dye was added to the second and fourth lane, respectively, to check whether further labeling can be achieved.

**(B)** Exemplary trace where cohesin core complex associated with a A550-labeled NIPBL-ΔN first extrudes downward. Upon exchange to a JF646-labeled NIPBL-ΔN at ~150 s, the cohesin complex extrudes upwards.

**(C)** Exemplary trace where a JF646-labeled NIPBL-ΔN resides at cohesin which extrudes solely unidirectionally (here towards the upper en(D).

**(D)** Example of DNA loop extrusion with exchange of NIPBL-ΔN without direction change.

**(E)** Example of DNA loop extrusion with exchange of NIPBL-ΔN and concomitant direction change between ~45 s and ~125 s. Another NIPBL-ΔN exchange occurs at ~240 s but the direction remains (downward here).

**(F)** Distribution of the time interval between NIPBL-ΔN disappearance and the end of the preceding extrusion phase (N = 45). The red bar represents the occurrences when NIPBL-ΔN disappearance coincided with the end of an extrusion event.

**(G)** Distribution of time between disappearance of one and appearance of the next NIPBL-ΔN molecule (N = 34).

**(H)** Diffusion constant of the DNA loop for diffusion phases in the presence (N = 45) and absence (N = 6) of NIPBL-ΔN. White dots denote median values, the box extends between first and third quartile, and whiskers extend to 1.5*IQR. Statistical significance was assessed by a Mann-Whitney test (not significant: $p > 0.05$).

**(I)** Loop shrinkage rate in the presence (N = 109) and absence (N = 11) of NIPBL-ΔN. White dots denote median values, the box extends between first and third quartile, and whiskers extend to 1.5*IQR. Statistical significance was assessed by a Mann-Whitney test (not significant: $p > 0.05$).

**(J)** Fraction of DNA molecules with a loop in DNA loop extrusion experiments using single-chain cohesin in 50 mM and 25 mM NaCl, with and without a 12-fold excess of STAG1 (N = 300, 1151, 1046, 878 from left to right). Dots represent the mean and error bars represent the SD of three independent experiments.

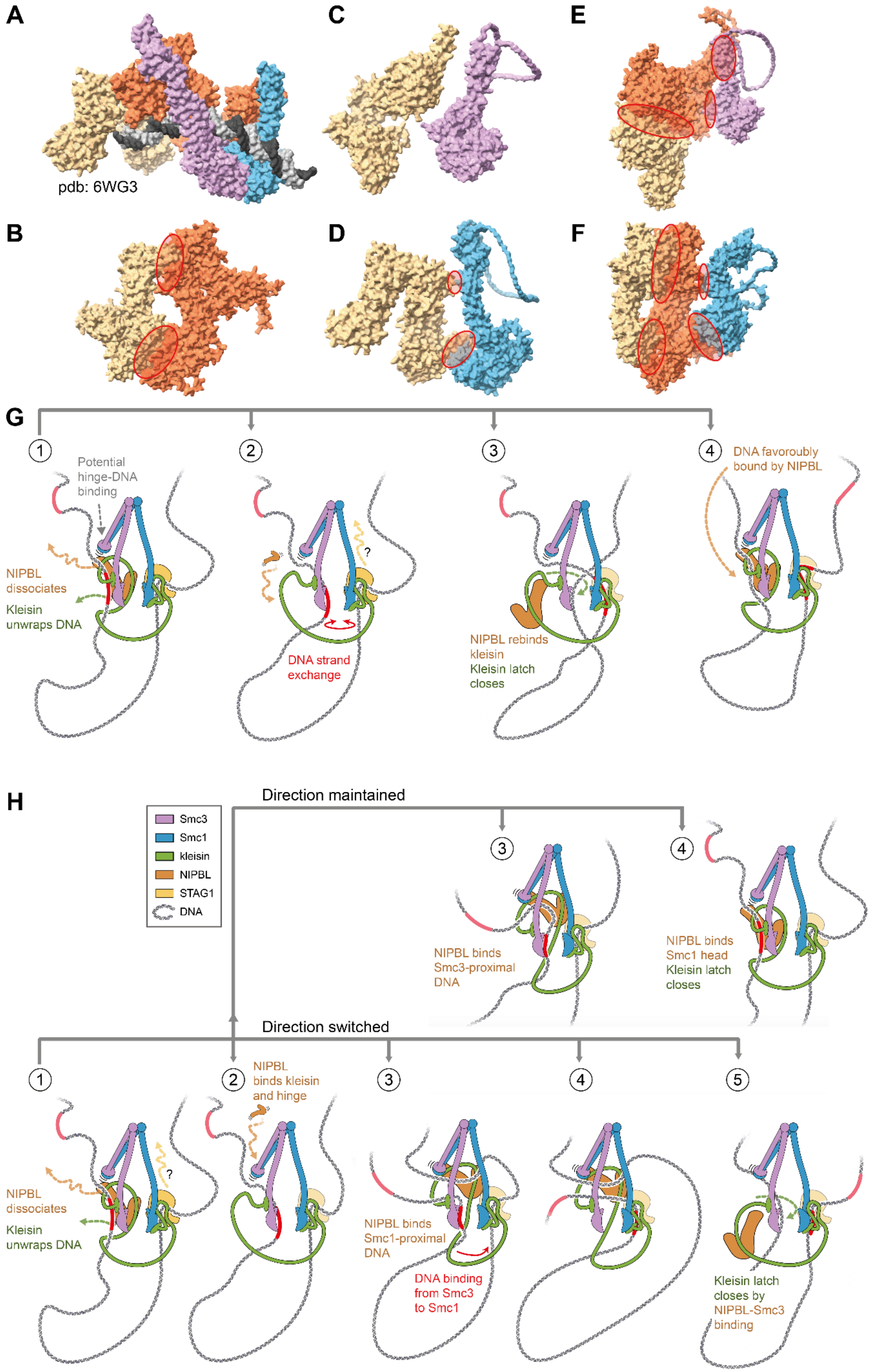
A
pdb: 6WG3
B
C
D
E
F
G
1
Potential hinge-DNA binding
NIPBL dissociates
Kleisin unwraps DNA
2
?
DNA strand exchange
3
NIPBL rebinds kleisin
Kleisin latch closes
4
DNA favouroubly bound by NIPBL
H
Smc3
Smc1
kleisin
NIPBL
STAG1
DNA
Direction maintained
3
NIPBL binds Smc3-proximal DNA
4
NIPBL binds Smc1 head
Kleisin latch closes
Direction switched
1
NIPBL dissociates
Kleisin unwraps DNA
?
2
NIPBL binds kleisin and hinge
3
NIPBL binds Smc1-proximal DNA
DNA binding from Smc3 to Smc1
4
5
Kleisin latch closes by NIPBL-Smc3 binding

**Figure S5: Potential pathways of loop extrusion direction change via DNA strand exchange mediated by NIPBL exchange. Related to Figure 5.**

**(A)** Experimentally solved structure of the cohesin-NIPBL-DNA complex in the engaged state (pdb 6WG3 , ref. [3]).

**(B)** AlphaFold2 prediction of the NIPBL-STAG1 heterodimer. Contact regions are indicated by red ellipses. Structure prediction yields similar results as in Nasmyth *et al.* (ref. [4]).

**(C)** AlphaFold2 prediction of the Smc3 head domain including a part of the coiled coils (connected by a 64x G linker) and STAG1. No interaction is apparent. Kleisin was included during the folding but is now shown for clarity.

**(D)** AlphaFold2 prediction of the Smc1 head domain including a part of the coiled coils (connected by a 64x G linker) and STAG1. Interactions between STAG1 and Smc1 are apparent close to DNA-binding patches 1 and 2 which were identified by Bauer *et al.* (ref. [5]). Kleisin was included during the folding but is now shown for clarity.

**(E)** AlphaFold2 prediction of the Smc3 head domain including a part of the coiled coils (connected by a 64x G linker), NIPBL, and STAG1. Interactions between Smc3 and NIPBL, as well as NIPBL and STAG1 are apparent, but not between Smc3 and STAG1. Kleisin was included during the folding but is now shown for clarity.

**(F)** AlphaFold2 prediction of the Smc1 head domain including a part of the coiled coils (connected by a 64x G linker), NIPBL, and STAG1. Interactions between Smc1 and NIPBL, as well as between NIPBL and STAG1 are apparent, but not between Smc1 and STAG1 directly. Kleisin was included during the folding but is now shown for clarity.

**(G)** Upon dissociation of NIPBL, the kleisin unwraps DNA (step 1). To prevent loss of the Smc3-proximal DNA (red), the Smc3 ATPase head or the Smc hinge may bind DNA. STAG1 may dissociate from cohesin concomitantly (step 2). DNA strand exchange may occur spontaneously between the ATPase heads (step 2). Upon rebinding of NIPBL to kleisin (step 3) and closing of the kleisin latch, the former Smc1-proximal strand is poised for the next loop extrusion cycle (step 4).

**(H)** Step 1 is as described in (G). Alternatively, to the pathway sketched in (A), NIPBL may re-bind the cohesin complex by binding to kleisin and the Smc hinge (step 2). The NIPBL-kleisin-hinge complex may bind either DNA strand which is the one on which loop extrusion subsequently proceeds. If the former Smc1-proximal DNA strand is chosen (step 3, lower panel), the Smc3-proximal DNA strand exchanges to the DNA binding site at the Smc1 ATPase head or at STAG1 (steps 3 and 4, lower panel). If the Smc3-proximal strand is chosen (step 3, upper panel), no direction change occurs. The binding of NIPBL-kleisin to the Smc3 ATPase head prepares the complex for the next loop extrusion cycle (step 5, lower panel; step 4, upper panel).

#### SUPPLEMENTARY REFERENCES


1. Taschner, M. & Gruber, S. DNA segment capture by Smc5/6 holocomplexes. *Nat Struct Mol Biol* **30**, 619–628 (2023).
2. Pradhan, B. *et al.* The Smc5/6 complex is a DNA loop-extruding motor. *Nature* **616**, 843–848 (2023).
3. Shi, Z., Gao, H., Bai, X. C. & Yu, H. Cryo-EM structure of the human cohesin-NIPBL-DNA complex. *Science* **368**, 1454–1459 (2020).
4. Nasmyth, K. A., Lee, B.-G., Roig, M. B. & Löwe, J. *What AlphaFold tells us about cohesin's retention on and release from chromosomes*. http://biorxiv.org/lookup/doi/10.1101/2023.04.14.536858 (2023) doi:10.1101/2023.04.14.536858.
5. Bauer, B. W. *et al.* Cohesin mediates DNA loop extrusion by a "swing and clamp" mechanism. *Cell* **184**, 5448–5464 (2021).

### SUPPLEMENTARY NOTE 1

The sequences used to generate AlphaFold2 predictions are listed below.

#### Smc3

GFLKLIEIENFKSYKGRQIIGPFQRFTAIIGPNGSGKSNLMDAISFVLGEKTSNLRVKTLRDLIHGAPVMGFLKLIEIENFKSYKGRQIIGPFQRFTAIIGPNGSGKSNLMDAISFVLGEKTSNLRVKTLRDLIHGAPV
GKPAANRAFVSMVYSEEGAEDRTFARVIVGGSSEYKINNKVVQLHEYSEELEKLGILIKARNFLVFQGAVESIAMKNPKERTALFEEISRSGELAQEYDKRKKEMVKAEEDTQFNYHRKKNIAAERKEAKQEKEE
ADRYQRLKDEVVRAQVQLQLFKLYHNEVEIEKLNKELASKNKEIEKDKKRMDKVEDELKEKKKELGKMMREQQQIEKEIKEKDSELNQKRPQYIKAKENTSHKIKKLEAAKKSLQNAQKHYKKRKGDMDELE
KEMLSVEKARQEFEERMEEESQSQGRDLTLEENQVKKYHRLKEEASKRAATLAQELEKFNRDQKADQDRLDLEERKKVETEAKIKQKLREIEENQKRIEKLEEYITTSKQSLEEQKKLEGELTEEVEMAKRRIDEI
NKELNQVMEQLGDARIDRQESSRQQRKAEIMESIKRLYPGSVYGRLIDLCQPTQKKYQIAVTKVLGKNMDAIIVDSEKTGRDCIQYIKEQRGEPETFLPLDYLEVKPTDEKLRELKGAKLVIDVIRYEPPHIKKAL
QYACGNALVCDNVEDARRIAFGGHQRHKTVALDGTLFQKSGVISGGASDLKAKARRWDEKAVDKLKEKKERLTEELKEQMKAKRKEAELRQVQSQAHGLQMRLKYSQSDLEQTKTRHLALNLQEKSKLES
ELANFGPRINDIKRIIQSREREMKDLKEKMNQVEDEVFEEFCREIGVRNIREFEEEKVKRQNEIAKKRLEFENQKTRLGIQLDFEKNQLKEDQDKVHMWEQTVKKDENEIEKLKKEEQRHMKIIDETMAQLQD
LKNQHLAKKSEVNDKNHEMEEIRKKLGGANKEMTHLQKEVTAIETKLEQKRSDRHNLLQACKMQDIKLPLSKGTMDDISQEEGSSQGEDSVSGSQRISSIYAREALIEIDYGDLCEDLKDAQAEEEIKQEMN
TLQQKLNEQQSVLQRIAAPNMKAMEKLESVRDKFQETSDEFEAARKRAKKAKQAFEQIKKERFDRFNACFESVATNIDEIYKALSRNSSAQAFLGPENPEEPYLDGINYNCVAPGKRFRPMDNLSGGEKTVA
ALALLFAIHSYKPAPFFVLDQIDAALDNTNIGKVANYIKEQSTCNFQAIVISLKEEFYTKAESLIGVYPEQGDCVISKVLTFDLTKYPDANPNPNEQ

#### Smc3 head complete (including 64G linker)

MGFLKLIEIENFKSYKGRQIIGPFQRFTAIIGPNGSGKSNLMDAISFVLGEKTSNLRVKTLRDLIHGAPVGKPAANRAFVSMVYSEEGAEDRTFARVIVGGSSEYKINNKVVQLHEYSEELEKLGILIKARNFLVFQ
GAVESIAMKNPKERTALFEEISRSGELAQEYDKRKKEMVKAEEDTQFNYHRKKNIAAERKEAKQGGGGGGGGGGGGGGGGGGGGGGGGGGGGGGGGGGGGGGGGGGGGGGGGGGGGGGGGGG
GGGGGLESVRDKFQETSDEFEAARKRAKKAKQAFEQIKKERFDRFNACFESVATNIDEIYKALSRNSSAQAFLGPENPEEPYLDGINYNCVAPGKRFRPMDNLSGGEKTVAALALLFAIHSYKPAPFFVLDQID
AALDNTNIGKVANYIKEQSTCNFQAIVISLKEEFYTKAESLIGVYPEQGDCVISKVLTFDLTKYPDANPNPNEQ

#### Smc1

MYIKQVIIQGFRSYRDQTIVDPFSSKHNVIVGRNGSGKSNFFYAIQFVLSDEFSHLRPEQRLALLHEGTGPRVISAFVEIIFDNSDNRLPIDKEEVSLRRVIGAKKDQYFLDKKMVTKNDVMNLLESAGFSRSNP
YYIVKQGKINQMATAPDSQRLKLLREVAGTRVYDERKEESISLMKETEGKREKINELLKYIEERLHTLEEEKEELAQYQKWDKMRRALEYTIYNQELNETRAKLDELSAKRETSGEKSRQLRDAQQDARDKMED
IERQVRELKTKISAMKEEKEQLSAERQEQIKQRTKLELKAKDLQDELAGNSEQRKRLLKERQKLLEKIEEKQKELAETEPKFNSVKEKEERGIARLAQATQERTDLYAKQGRGSQFTSKEERDKWIKKELKSLDQA
INDKKRQIAAIHKDLEDTEANKEKNLEQYNKLDQDLNEVKARVEELDRKYYEVKNKKDELQSERNYLWREENAEQQALAAKREDLEKKQQLLRAATGKAILNGIDSINKVLDHFRRKGINQHVQNGYHGIV
MNNFECEPAFYTCVEVTAGNRLFYHIVDSDEVSTKILMEFNKMNLPGEVTFLPLNKLDVRDTAYPETNDAIPMISKLRYNPRFDKAFKHVFGKTLICRSMEVSTQLARAFTMDCITLEGDQVSHRGALTGGYY
DTRKSRLELQKDVRKAEEELGELEAKLNENLRRNIERINNEIDQLMNQMQQIETQQRKFKASRDSILSEMKMLKEKRQQSEKTFMPKQRSLQSLEASLHAMESTRESLKAELGTDLLSQLSLEDQKRVDALN
DEIRQLQQENRQLLNERIKLEGIITRVETYLNENLRKRLDQVEQELNELRETEGGTVLTATTSELEAINKRVKDTMARSEDLDNSIDKTEAGIKELQKSMERWKNMEKEHMDAINHDTKELEKMTNRQGMLLK
KKEECMKKIRELGSLPQEAFEKYQTLSLKQLFRKLEQCNTELKKYSHVNKKALDQFVNFSEQKEKLIKRQEELDRGYKSIMELMNVLELRKYEAIQLTFKQVSKNFSEVFQKLVPGGKATLVMKKGDVEGSQSQ
DEGEGSGESERGSGSQSSVPSVDQFTGVGIRVSFTGKQGEMREMQQLSGGQKSLVALALIFAIQKCDPAPFYLFDQIDQALDAQHRKAVSDMIMELAVHAQFITTTFRPELLESADKFYGVKFRNKVSHIDVI
TAEMAKDFVEDDTTHG

#### Smc1 head complete (including 64G linker)

MYIKQVIIQGFRSYRDQTIVDPFSSKHNVIVGRNGSGKSNFFYAIQFVLSDEFSHLRPEQRLALLHEGTGPRVISAFVEIIFDNSDNRLPIDKEEVSLRRVIGAKKDQYFLDKKMVTKNDVMNLLESAGFSRSNP
YYIVKQGKINQMATAPDSQRLKLLREVAGTRVYDERKEESISLMKETEGKREKINELLKYIEERLHTLEEEKEELAQYQKWDKMRRALEYTIYNQELNETRAKLDEGGGGGGGGGGGGGGGGGGGGGGGGG
GGGGGGGGGGGGGGGGGGGGGGGGGGGGGGGGGGGGGGGNRQGMLLKKKEECMKKIRELGSLPQEAFEKYQTLSLKQLFRKLEQCNTELKKYSHVNKKALDQFVNFSEQKEKLIKRQEELDRGYKSI
MELMNVLELRKYEAIQLTFKQVSKNFSEVFQKLVPGGKATLVMKKGDVEGSQSQDEGEGSGESERGSGSQSSVPSVDQFTGVGIRVSFTGKQGEMREMQQLSGGQKSLVALALIFAIQKCDPAPFYLFDQI
DQALDAQHRKAVSDMIMELAVHAQFITTTFRPELLESADKFYGVKFRNKVSHIDVITAEMAKDFVEDDTTHG

#### Kleisin

MFYAHFVLSKRGPLAKIWLAAHWDKKLTKAHVFECNLESSVESIISPKVKMALRTSGHLLLGVVRIYHRKAKYLLADCNEAFIKIKMAFRPGVVDLPEENREAAYNAITLPEEFHDFDQPLPDLDDIDVAQQFS
LNQSRVEEITMREEVGNISILQENDFGDFGMDDREIMREGSAFEDDDMLVSTTTSNLLLESEQSTSNLNEKINHLEYEDQYKDDNFGEGNDGGILDDKLISNNDGGIFDDPPALSEAGVMLPEQPAHDDM
DEDDNVSMGGPDSPDSVDPVEPMPTMTDQTTLVPNEEEAFALEPIDITVKETKAKRKRKLIVDSVKELDSKTIRAQLSDYSDIVTTLDLAPPTKKLMMWKETGGVEKLFSLPAQPLWNNRLLKLFTRCLTPLV
PEDLRKRRKGGEADNLDEFLKEFENPEVPREDQQQQHQQRDVIDEPIIEEPSRLQESVMEASRTNIDESAMPPPPPQGVKRKAGQIDPEPVMPPQQVEQMEIPPVELPPEEPPNICQLIPELELLPEKEKEKEK
EKEDDEEEEDEDASGGDQDQEERRWNKRTQQMLHGLQRALAKTGAESISLLELCRNTNRKQAAAKFYSFLVLKKQQAIELTQEEPYSDIIATPGPRFHII

#### NIPBL

MMDSSTFKRFTASIENILDNLEDMDFTAFGDDDEIPQELLLGKHQLNELGSESAKIKAMGIMDKLSTDKTVKVLNILEKNIQDGSKLSTLLNHNNDTEEEERLWRDLIMERVTKSADACLTTINIMTSPNMPK
AVYIEDVIERVIQYTKFHLQNTLYPQYDPVYRLDPHGGGLLSSKAKRAKCSTHKQRVIVMLYNKVCDIVSSLSELLEIQLLTDTTILQVSSMGITPFFVENVSELQLCAIKLVTAVFSRYEKHRQLILEEIFTSLARLP
TSKRSLRNFRLNSSDMDGEPMYIQMVTALVLQLIQCVVHLPSSEKDSNAEEDSNKKIDQDVVITNSYETAMRTAQNFLSIFLKKCGSKQGEEDYRPLFENFVQDLLSTVNKPEWPAAELLLSLLGRLLVHQF
SNKSTEMALRVASLDYLGTVAARLRKDAVTSKMDQGSIERILKQVSGGEDEIQQLQKALLDYLDENTETDPSLVFSRKFYIAQWFRDTTLETEKAMKSQKDEESSEGTHHAKEIETTGQIMHRAENRKKFLRS
IIKTTPSQFSTLKMNSDTVDYDDACLIVRYLASMRPFAQSFDIYLTQILRVLGENAIAVRTKAMKCLSEVVAVDPSILARLDMQRGVHGRLMDNSTSVREAAVELLGRFVLCRPQLAEQYYDMLIERILDTGIS
VRKRVIKILRDICIEQPTFPKITEMCVKMIRRVNDEEGIKKLVNETFQKLWFTPTPHNDKEAMTRKILNITDVVAACRDTGYDWFEQLLQNLLKSEEDSSYKPVKKACTQLVDNLVEHILKYEESLADSDNKGV
NSGRLVACITTLFLFSKIRPQLMVKHAMTMQPYLTTKCSTQNDFMVICNVAKILELVVPLMEHPSETFLATIEEDLMKLIIKYGMTVVQHCVSCLGAVVNKVTQNFKFVWACFNRYYGAISKLKSQHQEDPN
NTSLLTNKPALLRSLFTVGALCRHFDFDLEDFKGNSKVNIKDKVLELLMYFTKHSDEEVQTKAIIGLGFAFIQHPSLMFEQEVKNLYNNILSDKNSSVNLKIQVLKNLQTYLQEEDTRMQQADRDWKKVAKQ
EDLKEMGDVSSGMSSSIMQLYLKQVLEAFFHTQSSVRHFALNVIALTLNQGLIHPVQCVPYLIAMGTDPEPAMRNKADQQLVEIDKKYAGFIHMKAVAGMKMSYQVQQAINTCLKDPVRGFRQDESSS
ALCSHLYSMIRGNRQHRRAFLISLLNLFDDTAKTDVTMLLYIADNLACFPYQTQEEPLFIMHHIDITLSVSGSNLLQSFKESMVKDKRKERKSSPSKENESSDSEEEVSRPRKSRKRVDSDSDSDSEDDINSVM
KCLPENSAPLIEFANVSQGILLLLMLKQHLKNLCGFSDSKIQKYSPSESAKVYDKAINRKTGVHFHPKQTLDFLRSDMANSKITEEVKRSIVKQYLDFKLLMEHL

#### STAG1

TLFEVVKLGKSAMQSVVDDWIESYKQDRDIALLDLINFFIQCSGCRGTVRIEMFRNMQNAEIIRKMTEEFDEDSGDYPLTMPGPQWKKFRSNFCEFIGVLIRQCQYSIIYDEYMMDTVISLLTGLSDSQVRAF
RHTSTLAAMKLMTALVNVALNLSIHQDNTQRQYEAERNKMIGKRANERLELLLQKRKELQENQDEIENMMNSIFKGIFVHRYRDAIAEIRAICIEEIGVWMKMYSDAFLNDSYLKYVGWTLHDRQGEVRL
KCLKALQSLYTNRELFPKLELFTNRFKDRIVSMTLDKEYDVAVEAIRLVTLILHGSEEALSNEDCENVYHLVYSAHRPVAVAAGEFLHKKLFSRHDPQAEEALAKRRGRNSPNGNLIRMLVLFFLESELHEHAAY
LVDSLWESSQELLKDWECMTELLLEEPVQGEEAMSDRQESALIELMVCTIRQAAEAHPPVGRGTGKRVLTAKERKTQIDDRNKLTEHFIITLPMLLSKYSADAEKVANLLQIPQYFDLEIYSTGRMEKHLDALL
KQIKFVVEKHVESDVLEACSKTYSILCSEEYTIQNRVDIARSQLIDEFVDRFNHSVEDLLQEGEEADDDDIYNVLSTLKRLTSFHNAHDLTKWDLFGNCYRLLKTGIEHGAMPEQIVVQALQCSHYSILWQLVK
ITDGSPSKEDLLVLRKTVKSFLAVCQQCLSNVNTPVKEQAFMLLCDLLMIFSHQLMTGGREGLQPLVFNPDTGLQSELLSFVMDHVFIDQDEENQSMEGDEEDEANKIEALHKRRNLLAAFSKLIIYDIVDM
HAAADIFKHYMKYYNDYGDIIKETLSKTRQIDKIQCAKTLILSLQQLFNELVQEQGPNLDRTSAHVSGIKELARRFALTFGLDQIKTREAVATLHKDGIEFAFKYQNQKGQEYPPPNLAFLEVLSEFSSKLLRQD
KKTVHSYLEKFLTEQMMERREDVWLPLISYRNSL

#### Smc3head:kleisin:SA1

MGFLKLIEIENFKSYKGRQIIGPFQRFTAIIGPNGSGKSNLMDAISFVLGEKTSNLRVKTLRDLIHGAPVGKPAANRAFVSMVYSEEGAEDRTFARVIVGGSSEYKINNKVVQLHEYSEELEKLGILIKARNFLVFQ
GAVESIAMKNPKERTALFEEISRSGELAQEYDKRKKEMVKAEEDTQFNYHRKKNIAAERKEAKQGGGGGGGGGGGGGGGGGGGGGGGGGGGGGGGGGGGGGGGGGGGGGGGGG
GGGGGLESVRDKFQETSDEFEAARKRAKKAKQAFEQIKKERFDRFNACFESVATNIDEIYKALSRNSSAQAFLGPENPEEPYLDGINYNCVAPGKRFRPMDNLSGGEKTVAALALLFAIHSYKPAPFFVLDQID
AALDNTNIGKVANYIKEQSTCNFQAIVISLKEEFYTKAESLIGVYPEQGDCVISKVLTFDLTKYPDANPNPNEQ:MFYAHFVLSKRGPLAKIWLAAHWDKKLTKAHVFECNLESSVESIISPKVKMALRTSGHLLL
GVVRIYHRKAKYLLADCNEAFIKIKMAFRPGVVDLPEENREAAYNAITLPEEFHDFDQPLPDLDDIDVAQQFSLNQSRVEEITMREEVGNISILQENDFGDFGMDDREIMREGSAFEDDDMLVSTTTSNLLLE
SEQSTSNLNEKINHLEYEDQYKDDNFGEGNDGGILDDKLISNNDGGIFDDPPALSEAGVMLPEQPAHDDMDEDDNVSMGGPDSPDSVDPVEPMPTMTDQTTLVPNEEEAFALEPIDITVKETKAKRKRKL
IVDSVKELDSKTIRAQLSDYSDIVTTLDLAPPTKKLMMWKETGGVEKLFSLPAQPLWNNRLLKLFTRCLTPLVPEDLRKRRKGGEADNLDEFLKEFENPEVPREDQQQQHQQRDVIDEPIIEEPSRLQESVMEA
SRTNIDESAMPPPPPQGVKRKAGQIDPEPVMPPQQVEQMEIPPVELPPEEPPNICQLIPELELLPEKEKEKEKEKEDDEEEEDEDASGGDQDQEERRWNKRTQQMLHGLQRALAKTGAESISLLELCRNTNR
KQAAAKFYSFLVLKKQQAIELTQEEPYSDIIATPGPRFHII:TLFEVVKLGKSAMQSVVDDWIESYKQDRDIALLDLINFFIQCSGCRGTVRIEMFRNMQNAEIIRKMTEEFDEDSGDYPLTMPGPQWKKFRSNF
CEFIGVLIRQCQYSIIYDEYMMDTVISLLTGLSDSQVRAFRHTSTLAAMKLMTALVNVALNLSIHQDNTQRQYEAERNKMIGKRANERLELLLQKRKELQENQDEIENMMNSIFKGIFVHRYRDAIAEIRAICIE
EIGVWMKMYSDAFLNDSYLKYVGWTLHDRQGEVRLKCLKALQSLYTNRELFPKLELFTNRFKDRIVSMTLDKEYDVAVEAIRLVTLILHGSEEALSNEDCENVYHLVYSAHRPVAVAAGEFLHKKLFSRHDPQ
AEEALAKRRGRNSPNGNLIRMLVLFFLESELHEHAAYLVDSLWESSQELLKDWECMTELLLEEPVQGEEAMSDRQESALIELMVCTIRQAAEAHPPVGRGTGKRVLTAKERKTQIDDRNKLTEHFIITLPMLLSK
YSADAEKVANLLQIPQYFDLEIYSTGRMEKHLDALLKQIKFVVEKHVESDVLEACSKTYSILCSEEYTIQNRVDIARSQLIDEFVDRFNHSVEDLLQEGEEADDDDIYNVLSTLKRLTSFHNAHDLTKWDLFGNC
YRLLKTGIEHGAMPEQIVVQALQCSHYSILWQLVKITDGSPSKEDLLVLRKTVKSFLAVCQQCLSNVNTPVKEQAFMLLCDLLMIFSHQLMTGGREGLQPLVFNPDTGLQSELLSFVMDHVFIDQDEENQSM
EGDEEDEANKIEALHKRRNLLAAFSKLIIYDIVDMHAAADIFKHYMKYYNDYGDIIKETLSKTRQIDKIQCAKTLILSLQQLFNELVQEQGPNLDRTSAHVSGIKELARRFALTFGLDQIKTREAVATLHKDGIEFA
FKYQNQKGQEYPPPNLAFLEVLSEFSSKLLRQDKKTVHSYLEKFLTEQMMERREDVWLPLISYRNSL

#### Smc1head:kleisin:SA1

MYIKQVIIQGFRSYRDQTIVDPFSSKHNVIVGRNGSGKSNFFYAIQFVLSDEFSHLRPEQRLALLHEGTGPRVISAFVEIIFDNSDNRLPIDKEEVSLRRVIGAKKDQYFLDKKMVTKNDVMNLLESAGFSRSNP
YYIVKQGKINQMATAPDSQRLKLLREVAGTRVYDERKEESISLMKETEGKREKINELLKYIEERLHTLEEEKEELAQYQKWDKMRRALEYTIYNQELNETRAKLDEGGGGGGGGGGGGGGGG
GGGGGGGGGGGGGGGGGGGGGGGGGGGGGGGGGGGGGNRQGMLLKKKEECMKKIRELGSLPQEAFEKYQTLSLKQLFRKLEQCNTELKKYSHVNKKALDQFVNFSEQKEKLIKRQEELDRGYKSI
MELMNVLELRKYEAIQLTFKQVSKNFSEVFQKLVPGGKATLVMKKGDVEGSQSQDEGEGSGESERGSGSQSSVPSVDQFTGVGIRVSFTGKQGEMREMQQLSGGQKSLVALALIFAIQKCDPAPFYLFDQID
QALDAQHRKAVSDMIMELAVHAQFITTTFRPELLESADKFYGVKFRNKVSHIDVITAEMAKDFVEDDTTHG:MFYAHFVLSKRGPLAKIWLAAHWDKKLTKAHVFECNLESSVESIISPKVKMALRTSGHLLL
GVVRIYHRKAKYLLADCNEAFIKIKMAFRPGVVDLPEENREAAYNAITLPEEFHDFDQPLPDLDDIDVAQQFSLNQSRVEEITMREEVGNISILQENDFGDFGMDDREIMREGSAFEDDDMLVSTTTSNLLLE
SEQSTSNLNEKINHLEYEDQYKDDNFGEGNDGGILDDKLISNNDGGIFDDPPALSEAGVMLPEQPAHDDMDEDDNVSMGGPDSPDSVDPVEPMPTMTDQTTLVPNEEEAFALEPIDITVKETKAKRKRKL
IVDSVKELDSKTIRAQLSDYSDIVTTLDLAPPTKKLMMWKETGGVEKLFSLPAQPLWNNRLLKLFTRCLTPLVPEDLRKRRKGGEADNLDEFLKEFENPEVPREDQQQQHQQRDVIDEPIIEEPSRLQESVMEA
SRTNIDESAMPPPPPQGVKRKAGQIDPEPVMPPQQVEQMEIPPVELPPEEPPNICQLIPELELLPEKEKEKEKEKEDDEEEEDEDASGGDQDQEERRWNKRTQQMLHGLQRALAKTGAESISLLELCRNTNR
KQAAAKFYSFLVLKKQQAIELTQEEPYSDIIATPGPRFHII:TLFEVVKLGKSAMQSVVDDWIESYKQDRDIALLDLINFFIQCSGCRGTVRIEMFRNMQNAEIIRKMTEEFDEDSGDYPLTMPGPQWKKFRSNF
CEFIGVLIRQCQYSIIYDEYMMDTVISLLTGLSDSQVRAFRHTSTLAAMKLMTALVNVALNLSIHQDNTQRQYEAERNKMIGKRANERLELLLQKRKELQENQDEIENMMNSIFKGIFVHRYRDAIAEIRAICIE
EIGVWMKMYSDAFLNDSYLKYVGWTLHDRQGEVRLKCLKALQSLYTNRELFPKLELFTNRFKDRIVSMTLDKEYDVAVEAIRLVTLILHGSEEALSNEDCENVYHLVYSAHRPVAVAAGEFLHKKLFSRHDPQ
AEEALAKRRGRNSPNGNLIRMLVLFFLESELHEHAAYLVDSLWESSQELLKDWECMTELLLEEPVQGEEAMSDRQESALIELMVCTIRQAAEAHPPVGRGTGKRVLTAKERKTQIDDRNKLTEHFIITLPMLLSK
YSADAEKVANLLQIPQYFDLEIYSTGRMEKHLDALLKQIKFVVEKHVESDVLEACSKTYSILCSEEYTIQNRVDIARSQLIDEFVDRFNHSVEDLLQEGEEADDDDIYNVLSTLKRLTSFHNAHDLTKWDLFGNC
YRLLKTGIEHGAMPEQIVVQALQCSHYSILWQLVKITDGSPSKEDLLVLRKTVKSFLAVCQQCLSNVNTPVKEQAFMLLCDLLMIFSHQLMTGGREGLQPLVFNPDTGLQSELLSFVMDHVFIDQDEENQSM
EGDEEDEANKIEALHKRRNLLAAFSKLIIYDIVDMHAAADIFKHYMKYYNDYGDIIKETLSKTRQIDKIQCAKTLILSLQQLFNELVQEQGPNLDRTSAHVSGIKELARRFALTFGLDQIKTREAVATLHKDGIEFA
FKYQNQKGQEYPPPNLAFLEVLSEFSSKLLRQDKKTVHSYLEKFLTEQMMERREDVWLPLISYRNSL

#### Smc3head:kleisin:NIPBL:SA1

MGFLKLIEIENFKSYKGRQIIGPFQRFTAIIGPNGSGKSNLMDAISFVLGEKTSNLRVKTLRDLIHGAPVGKPAANRAFVSMVYSEEGAEDRTFARVIVGGSSEYKINNKVVQLHEYSEELEKLGILIKARNFLVFQ
GAVESIAMKNPKERTALFEEISRSGELAQEYDKRKKEMVKAEEDTQFNYHRKKNIAAERKEAKQGGGGGGGGGGGGGGGGGGGGGGGGGGGGGGGGGGGGGGGGGGGGGGGGG
GGGGGGLESVRDKFQETSDEFEAARKRAKKAKQAFEQIKKERFDRFNACFESVATNIDEIYKALSRNSSAQAFLGPENPEEPYLDGINYNCVAPGKRFRPMDNLSGGEKTVAALALLFAIHSYKPAPFFVLDQ
IDAALDNTNIGKVANYIKEQSTCNFQAIVISLKEEFYTKAESLIGVYPEQGDCVISKVLTFDLTKYPDANPNPNEQ:MFYAHFVLSKRGPLAKIWLAAHWDKKLTKAHVFECNLESSVESIISPKVKMALRTSGH
LLLGVVRIYHRKAKYLLADCNEAFIKIKMAFRPGVVDLPEENREAAYNAITLPEEFHDFDQPLPDLDDIDVAQQFSLNQSRVEEITMREEVGNISILQENDFGDFGMDDREIMREGSAFEDDDMLVSTTTSNL
LLESEQSTSNLNEKINHLEYEDQYKDDNFGEGNDGGILDDKLISNNDGGIFDDPPALSEAGVMLPEQPAHDDMDEDDNVSMGGPDSPDSVDPVEPMPTMTDQTTLVPNEEEAFALEPIDITVKETKAKRK
RKLIVDSVKELDSKTIRAQLSDYSDIVTTLDLAPPTKKLMMWKETGGVEKLFSLPAQPLWNNRLLKLFTRCLTPLVPEDLRKRRKGGEADNLDEFLKEFENPEVPREDQQQQHQQRDVIDEPIIEEPSRLQESV
MEASRTNIDESAMPPPPPQGVKRKAGQIDPEPVMPPQQVEQMEIPPVELPPEEPPNICQLIPELELLPEKEKEKEKEKEDDEEEEDEDASGGDQDQEERRWNKRTQQMLHGLQRALAKTGAESISLLELCRN
TNRKQAAAKFYSFLVLKKQQAIELTQEEPYSDIIATPGPRFHII:MMDSSTFKRFTASIENILDNLEDMDFTAFGDDDEIPQELLLGKHQLNELGSESAKIKAMGIMDKLSTDKTVKVLNILEKNIQDGSKLSTLL
NHNNDTEEEERLWRDLIMERVTKSADACLTTINIMTSPNMPKAVYIEDVIERVIQYTKFHLQNTLYPQYDPVYRLDPHGGGLLSSKAKRAKCSTHKQRVIVMLYNKVCDIVSSLSELLEIQLLTDTTILQVSSM
GITPFFVENVSELQLCAIKLVTAVFSRYEKHRQLILEEIFTSLARLPTSKRSLRNFRLNSSDMDGEPMYIQMVTALVLQLIQCVVHLPSSEKDSNAEEDSNKKIDQDVVITNSYETAMRTAQNFLSIFLKKCGSKQ
GEEDYRPLFENFVQDLLSTVNKPEWPAAELLLSLLGRLLVHQFSNKSTEMALRVASLDYLGTVAARLRKDAVTSKMDQGSIERILKQVSGGEDEIQQLQKALLDYLDENTETDPSLVFSRKFYIAQWFRDTTL
ETEKAMKSQKDEESSEGTHHAKEIETTGQIMHRAENRKKFLRSIIKTTPSQFSTLKMNSDTVDYDDACLIVRYLASMRPFAQSFDIYLTQILRVLGENAIAVRTKAMKCLSEVVAVDPSILARLDMQRGVHGR
LMDNSTSVREAAVELLGRFVLCRPQLAEQYYDMLIERILDTGISVRKRVIKILRDICIEQPTFPKITEMCVKMIRRVNDEEGIKKLVNETFQKLWFTPTPHNDKEAMTRKILNITDVVAACRDTGYDWFEQLLQ
NLLKSEEDSSYKPVKKACTQLVDNLVEHILKYEESLADSDNKGVNSGRLVACITTLFLFSKIRPQLMVKHAMTMQPYLTTKCSTQNDFMVICNVAKILELVVPLMEHPSETFLATIEEDLMKLIIKYGMTVVQH
CVSCLGAVVNKVTQNFKFVWACFNRYYGAISKLKSQHQEDPNNTSLLTNKPALLRSLFTVGALCRHFDFDLEDFKGNSKVNIKDKVLELLMYFTKHSDEEVQTKAIIGLGFAFIQHPSLMFEQEVKNLYNNIL
SDKNSSVNLKIQVLKNLQTYLQEEDTRMQQADRDWKKVAKQEDLKEMGDVSSGMSSSIMQLYLKQVLEAFFHTQSSVRHFALNVIALTLNQGLIHPVQCVPYLIAMGTDPEPAMRNKADQQLVEIDKK
YAGFIHMKAVAGMKMSYQVQQAINTCLKDPVRGFRQDESSSALCSHLYSMIRGNRQHRRAFLISLLNLFDDTAKTDVTMLLYIADNLACFPYQTQEEPLFIMHHIDITLSVSGSNLLQSFKESMVKDKRKE
RKSSPSKENESSDSEEEVSRPRKSRKRVDSDSDSDSEDDINSVMKCLPENSAPLIEFANVSQGILLLLMLKQHLKNLCGFSDSKIQKYSPSESAKVYDKAINRKTGVHFHPKQTLDFLRSDMANSKITEEVKRSI
VKQYLDFKLLMEHL:TLFEVVKLGKSAMQSVVDDWIESYKQDRDIALLDLINFFIQCSGCRGTVRIEMFRNMQNAEIIRKMTEEFDEDSGDYPLTMPGPQWKKFRSNFCEFIGVLIRQCQYSIIYDEYMMDTV
ISLLTGLSDSQVRAFRHTSTLAAMKLMTALVNVALNLSIHQDNTQRQYEAERNKMIGKRANERLELLLQKRKELQENQDEIENMMNSIFKGIFVHRYRDAIAEIRAICIEEIGVWMKMYSDAFLNDSYLKYV
GWTLHDRQGEVRLKCLKALQSLYTNRELFPKLELFTNRFKDRIVSMTLDKEYDVAVEAIRLVTLILHGSEEALSNEDCENVYHLVYSAHRPVAVAAGEFLHKKLFSRHDPQAEEALAKRRGRNSPNGNLIRML
VLFFLESELHEHAAYLVDSLWESSQELLKDWECMTELLLEEPVQGEEAMSDRQESALIELMVCTIRQAAEAHPPVGRGTGKRVLTAKERKTQIDDRNKLTEHFIITLPMLLSKYSADAEKVANLLQIPQYFDLEI
YSTGRMEKHLDALLKQIKFVVEKHVESDVLEACSKTYSILCSEEYTIQNRVDIARSQLIDEFVDRFNHSVEDLLQEGEEADDDDIYNVLSTLKRLTSFHNAHDLTKWDLFGNCYRLLKTGIEHGAMPEQIVVQA
LQCSHYSILWQLVKITDGSPSKEDLLVLRKTVKSFLAVCQQCLSNVNTPVKEQAFMLLCDLLMIFSHQLMTGGREGLQPLVFNPDTGLQSELLSFVMDHVFIDQDEENQSMEGDEEDEANKIEALHKRRNL
LAAFSKLIIYDIVDMHAAADIFKHYMKYYNDYGDIIKETLSKTRQIDKIQCAKTLILSLQQLFNELVQEQGPNLDRTSAHVSGIKELARRFALTFGLDQIKTREAVATLHKDGIEFAFKYQNQKGQEYPPPNLAFL
EVLSEFSSKLLRQDKKTVHSYLEKFLTEQMMERREDVWLPLISYRNSL

#### Smc1head:kleisin:NIPBL:SA1

MYIKQVIIQGFRSYRDQTIVDPFSSKHNVIVGRNGSGKSNFFYAIQFVLSDEFSHLRPEQRLALLHEGTGPRVISAFVEIIFDNSDNRLPIDKEEVSLRRVIGAKKDQYFLDKKMVTKNDVMNLLESAGFSRSNP
YYIVKQGKINQMATAPDSQRLKLLREVAGTRVYDERKEESISLMKETEGKREKINELLKYIEERLHTLEEEKEELAQYQKWDKMRRALEYTIYNQELNETRAKLDEGGGGGGGGGGGGGGGG
GGGGGGGGGGGGGGGGGGGGGGGGGGGGGGGGGGGGGNRQGMLLKKKEECMKKIRELGSLPQEAFEKYQTLSLKQLFRKLEQCNTELKKYSHVNKKALDQFVNFSEQKEKLIKRQEELDRGYKSI
MELMNVLELRKYEAIQLTFKQVSKNFSEVFQKLVPGGKATLVMKKGDVEGSQSQDEGEGSGESERGSGSQSSVPSVDQFTGVGIRVSFTGKQGEMREMQQLSGGQKSLVALALIFAIQKCDPAPFYLFDQI
DQALDAQHRKAVSDMIMELAVHAQFITTTFRPELLESADKFYGVKFRNKVSHIDVITAEMAKDFVEDDTTHG:MFYAHFVLSKRGPLAKIWLAAHWDKKLTKAHVFECNLESSVESIISPKVKMALRTSGHL
LLGVVRIYHRKAKYLLADCNEAFIKIKMAFRPGVVDLPEENREAAYNAITLPEEFHDFDQPLPDLDDIDVAQQFSLNQSRVEEITMREEVGNISILQENDFGDFGMDDREIMREGSAFEDDDMLVSTTTSNLL
LESEQSTSNLNEKINHLEYEDQYKDDNFGEGNDGGILDDKLISNNDGGIFDDPPALSEAGVMLPEQPAHDDMDEDDNVSMGGPDSPDSVDPVEPMPTMTDQTTLVPNEEEAFALEPIDITVKETKAKRK
RKLIVDSVKELDSKTIRAQLSDYSDIVTTLDLAPPTKKLMMWKETGGVEKLFSLPAQPLWNNRLLKLFTRCLTPLVPEDLRKRRKGGEADNLDEFLKEFENPEVPREDQQQQHQQRDVIDEPIIEEPSRLQESV
MEASRTNIDESAMPPPPPQGVKRKAGQIDPEPVMPPQQVEQMEIPPVELPPEEPPNICQLIPELELLPEKEKEKEKEKEDDEEEEDEDASGGDQDQEERRWNKRTQQMLHGLQRALAKTGAESISLLELCRN
TNRKQAAAKFYSFLVLKKQQAIELTQEEPYSDIIATPGPRFHII:MMDSSTFKRFTASIENILDNLEDMDFTAFGDDDEIPQELLLGKHQLNELGSESAKIKAMGIMDKLSTDKTVKVLNILEKNIQDGSKLSTLL
NHNNDTEEEERLWRDLIMERVTKSADACLTTINIMTSPNMPKAVYIEDVIERVIQYTKFHLQNTLYPQYDPVYRLDPHGGGLLSSKAKRAKCSTHKQRVIVMLYNKVCDIVSSLSELLEIQLLTDTTILQVSSM

GITPFFVENVSELQLCAIKLVTAVFSRYEKHRQLILEEIFTSLARLPTSKRSLRNFRLNSSDMDGEPMYIQMVTALVLQLIQCVVHLPSSEKDSNAEEDSNKKIDQDVVITNSYETAMRTAQNFLSIFLKKCGSKQ
GEEDYRPLFENFVQDLLSTVNKPEWPAAELLLSLLGRLLVHQFSNKSTEMALRVASLDYLGTVAARLRKDAVTSKMDQGSIERILKQVSGGEDEIQQLQKALLDYLDENTETDPSLVFSRKFYIAQWFRDTTL
ETEKAMKSQKDEESSEGTHHAKEIETTGQIMHRAENRKKFLRSIIKTTPSQFSTLKMNSDTVDYDDACLIVRYLASMRPFAQSFDIYLTQILRVLGENAIAVRTKAMKCLSEVVAVDPSILARLDMQRGVHGR
LMDNSTSVREAAVELLGRFVLCRPQLAEQYYDMLIERILDTGISVRKRVIKILRDICIEQPTFPKITEMCVKMIRRVNDEEGIKKLVNETFQKLWFTPTPHNDKEAMTRKILNITDVVAACRDTGYDWFEQLLQ
NLLKSEEDSSYKPVKKACTQLVDNLVEHILKYEESLADSDNKGVNSGRLVACITTLFLFSKIRPQLMVKHAMTMQPYLTTKCSTQNDFMVICNVAKILELVVPLMEHPSETFLATIEEDLMKLIIKYGMTVVQH
CVSCLGAVVNKVTQNFKFVWACFNRYYGAISKLKSQHQEDPNNTSLLTNKPALLRSLFTVGALCRHFDFDLEDFKGNSKVNIKDKVLELLMYFTKHSDEEVQTKAIIGLGFAFIQHPSLMFEQEVKNLYNNIL
SDKNSSVNLKIQVLKNLQTYLQEEDTRMQQADRDWKKVAKQEDLKEMGDVSSGMSSSIMQLYLKQVLEAFFHTQSSVRHFALNVIALTLNQGLIHPVQCVPYLIAMGTDPEPAMRNKADQQLVEIDKK
YAGFIHMKAVAGMKMSYQVQQAINTCLKDPVRGFRQDESSSALCSHLYSMIRGNRQHRRAFLISLLNLFDDTAKTDVTMLLYIADNLACFPYQTQEEPLFIMHHIDITLSVSGSNLLQSFKESMVKDKRKE
RKSSPSKENESSDSEEEVSRPRKSRKRVDSDSDSDSEDDINSVMKCLPENSAPLIEFANVSQGILLLLMLKQHLKNLCGFSDSKIQKYSPSESAKVYDKAINRKTGVHFHPKQTLDFLRSDMANSKITEEVKRSI
VKQYLDFKLLMEHL:TLFEVVKLGKSAMQSVVDDWIESYKQDRDIALLDLINFFIQCSGCRGTVRIEMFRNMQNAEIIRKMTEEFDEDSGDYPLTMPGPQWKKFRSNFCEFIGVLIRQCQYSIIYDEYMMDTV
ISLLTGLSDSQVRAFRHTSTLAAMKLMTALVNVALNLSIHQDNTQRQYEAERNKMIGKRANERLELLLQKRKELQENQDEIENMMNSIFKGIFVHRYRDAIAEIRAICIEEIGVWMKMYSDAFLNDSYLKYV
GWTLHDRQGEVRLKCLKALQSLYTNRELFPKLELFTNRFKDRIVSMTLDKEYDVAVEAIRLVTLILHGSEEALSNEDCENVYHLVYSAHRPVAVAAGEFLHKKLFSRHDPQAEEALAKRRGRNSPNGNLIRML
VLFFLESELHEHAAYLVDSLWESSQELLKDWECMTELLLEEPVQGEEAMSDRQESALIELMVCTIRQAAEAHPPVGRGTGKRVLTAKERKTQIDDRNKLTEHFIITLPMLLSKYSADAEKVANLLQIPQYFDLEI
YSTGRMEKHLDALLKQIKFVVEKHVESDVLEACSKTYSILCSEEYTIQNRVDIARSQLIDEFVDRFNHSVEDLLQEGEEADDDDIYNVLSTLKRLTSFHNAHDLTKWDLFGNCYRLLKTGIEHGAMPEQIVVQA
LQCSHYSILWQLVKITDGSPSKEDLLVLRKTVKSFLAVCQQCLSNVNTPVKEQAFMLLCDLLMIFSHQLMTGGREGLQPLVFNPDTGLQSELLSFVMDHVFIDQDEENQSMEGDEEDEANKIEALHKRRNL
LAAFSKLIIYDIVDMHAAADIFKHYMKYYNDYGDIIKETLSKTRQIDKIQCAKTLILSLQQLFNELVQEQGPNLDRTSAHVSGIKELARRFALTFGLDQIKTREAVATLHKDGIEFAFKYQNQKGQEYPPPNLAFL
EVLSEFSSKLLRQDKKTVHSYLEKFLTEQMMERREDVWLPLISYRNSL
